# Beta-cell specific *Insr* deletion promotes insulin hypersecretion and improves glucose tolerance prior to global insulin resistance

**DOI:** 10.1101/2020.10.15.338160

**Authors:** Søs Skovsø, Evgeniy Panzhinskiy, Jelena Kolic, Haoning Howard Cen, Derek A. Dionne, Xiao-Qing Dai, Rohit B. Sharma, Lynda Elghazi, Cara E. Ellis, Katharine Faulkner, Stephanie A.M. Marcil, Peter Overby, Nilou Noursadeghi, Daria Hutchinson, Xiaoke Hu, Hong Li, Honey Modi, Jennifer S. Wildi, J. Diego Botezelli, Hye Lim Noh, Sujin Suk, Brian Gablaski, Austin Bautista, Ryekjang Kim, Corentin Cras-Méneur, Stephane Flibotte, Sunita Sinha, Dan S. Luciani, Corey Nislow, Elizabeth J. Rideout, Eric N. Cytrynbaum, Jason K. Kim, Ernesto Bernal-Mizrachi, Laura C. Alonso, Patrick E. MacDonald, James D. Johnson

## Abstract

Abstract

Insulin receptor (Insr) protein can be found at higher levels in pancreatic β-cells than in most other tissues, but the consequences of β-cell insulin resistance remain enigmatic. *Ins1*^cre^ allele was used to delete *Insr* specifically in β-cells of both female and male mice. Experimental mice were compared to *Ins1*^cre^-containing littermate controls at multiple ages and on multiple diets. RNA-seq of purified recombined β-cells revealed transcriptomic consequences of *Insr* loss, which differed between female and male mice. Action potential and calcium oscillation frequencies were increased in *Insr* knockout β- cells from female, but not male mice, whereas only male β*Insr*^KO^ mice had reduced ATP-coupled oxygen consumption rate and reduced expression of genes involved in ATP synthesis. Female β*Insr*^KO^ and β*Insr*^HET^ mice exhibited elevated insulin release in perifusion experiments, during hyperglycemic clamps, and following *i.p.* glucose challenge. Deletion of *Insr* did not alter β-cell area up to 9 months of age, nor did it impair hyperglycemia-induced proliferation. Based on our data, we adapted a mathematical model to include β-cell insulin resistance, which predicted that β-cell *Insr* knockout would improve glucose tolerance depending on the degree of whole-body insulin resistance. Indeed, glucose tolerance was significantly improved in female β*Insr*^KO^ and β*Insr*^HET^ mice when compared to controls at 9, 21 and 39 weeks, and also in insulin-sensitive 4-week old males. We did not observe improved glucose tolerance in older male mice or in high fat diet-fed mice, corroborating the prediction that global insulin resistance obscures the effects of β-cell specific insulin resistance. The propensity for hyperinsulinemia was associated with mildly reduced fasting glucose and increased body weight. We further validated our main *in vivo* findings using the *Ins1*-CreERT transgenic line and found that male mice had improved glucose tolerance 4 weeks after tamoxifen-mediated *Insr* deletion. Collectively, our data show that loss of β-cell *Insr* contributes to glucose-induced hyperinsulinemia, thereby improving glucose homeostasis in otherwise insulin sensitive dietary and age contexts.

## Introduction

Type 2 diabetes is a multifactorial disease. Several cell types, most prominently pancreatic β-cells, are dysfunctional prior to and after diagnosis^1^. Hyperinsulinemia, insulin resistance, impaired fasting glucose, and impaired glucose tolerance can all be observed prior to the onset of frank diabetes^2^, but the causal relationships between these factors remain incompletely understood^3^. Impaired insulin receptor (Insr) signaling is associated with obesity and often precedes the onset of overt type 2 diabetes, but it has been studied primarily in skeletal muscle, fat, and liver where it manifests differently^4^. Recent work in mice has established that β-cell specific insulin resistance can be observed early in the progression towards type 2 diabetes, when hyperinsulinemia is prominent, and independently of insulin resistance in other tissues^5^. The physiological consequences of reduced Insr in β-cells remain controversial.

It remains unresolved whether physiological insulin action on β-cells manifests as positive feedback to stimulate further insulin secretion, or negative feedback to inhibit its own release^6^. Human studies provide evidence for both possibilities. *In vivo* hyperinsulinemic-euglycemic clamps can reduce circulating C-peptide, a marker of endogenous insulin secretion^7, 8^. In some studies, this inhibition was impaired in the obese state suggesting that systemic insulin resistance also extends to the β-cells^8^. Bouche and colleagues replicated this result at basal glucose, but also found evidence that insulin can potentiate glucose-stimulated insulin secretion under specific conditions^9^. Others have shown that insulin can either stimulate or inhibit its own secretion depending on the metabolic context^10^. Administration of insulin to single β-cells *in vitro* increases intracellular calcium (Ca^2+^)^11^ and, in some studies, stimulates exocytosis^12^. However, Ca^2+^ release from intracellular stores is not always sufficient to evoke insulin exocytosis. Studies in human β-cells did not detect robust exocytosis or C-peptide release in response to exogenous insulin despite observed changes in Ca^2+^ release^13^.

Whether chronic deviations in autocrine insulin signaling affect β-cell development, survival and adaptation conditions is also controversial. Mice with chronically reduced insulin production have impaired β-cell expansion in the context of a high fat diet^14^. *In vitro*, physiologically relevant concentrations of insulin support the survival of both human and mouse β-cells^15^. We also reported insulin is sufficient to increase proliferation of cultured primary mouse β-cells and that blocking insulin secretion with somatostatin blunts proliferation induced by hyperglycemia^16^ and that the majority of glucose-dependent changes in gene expression in MIN6 cells are Insr-dependent^17^. However, hyperglycemia-induced β-cell proliferation has been proposed to bypass Insr^18, 19^. Thus, whether these signals from insulin and/or glucose are transmitted through Insr, Igf1r, or both receptors, remains unresolved.

To address the short- and long-term consequences of eliminating Insr signaling *in vivo*, Kulkarni and colleagues crossed mice with floxed *Insr* alleles and an *Ins2* promoter driven Cre transgene^20^. Using this and related models, they reported that mice lacking β-cell *Insr* had profound glucose intolerance and frank diabetes in some cases, due to impaired glucose-stimulated insulin secretion, Glut2 loss, and insufficient β-cell mass^20–22^. Their *Insr* deficient mice failed to exhibit the compensatory increase in β- cell mass that accompanies a high fat diet^21^. Doubt was cast on these results when these Cre lines were subsequently shown to have off-target tissue effects owing to endogenous *Ins2* expression in the brain and thymus^14, 23–26^. More recently, Wang and colleagues studied the roles of β-cell *Insr in utero* and in adult mice using an inducible *Ins1*-CreER transgenic mouse model^27, 28^, but these studies are confounded by the presence of the human growth hormone (hGH) minigene^29^, which necessitates the use of Cre-containing controls exclusively.

In the present study, we primarily used the constitutive *Ins1*^Cre^ knock-in strain with robust and specific recombination in β-cells^30^ to precisely reduce Insr signaling and define its consequences on glucose homeostasis. We validated our findings on glucose homeostasis using an additional β-cell specific model. Using this approach, we find clear evidence that Insr signaling plays a suppressive role on insulin secretion by modulating β-cell electrical excitability and that this effect is absent in conditions of global insulin resistance.

## Results

### Insr abundance in human islets and β-cells

We initiated our studies by conducting an unbiased analysis of insulin receptor abundance across tissues using publicly accessible data. Pancreatic islets had the 2^nd^ highest protein abundance of both isoforms of the insulin receptor across a panel of 24 human tissues, as quantified by mass-spectrometry (Fig. 1A). These results, which are not complicated by the limitations associated with anti-Insr antibodies, show that human islets can have more INSR protein than ‘classical’ insulin target tissues, including the liver and adipose. This also supports our previous observations suggesting that insulin receptors are more abundant in β-cells relative to neighbouring cells in the mouse pancreas^31^. Compilation of open source public single-cell RNA sequencing data from human islets demonstrated insulin receptor mRNA in 62.4% β-cells, alongside other islet cell types (Fig. 1A,B). Clearly, β-cells have evolved for Insr-mediated autocrine signaling.

**Figure 1.**
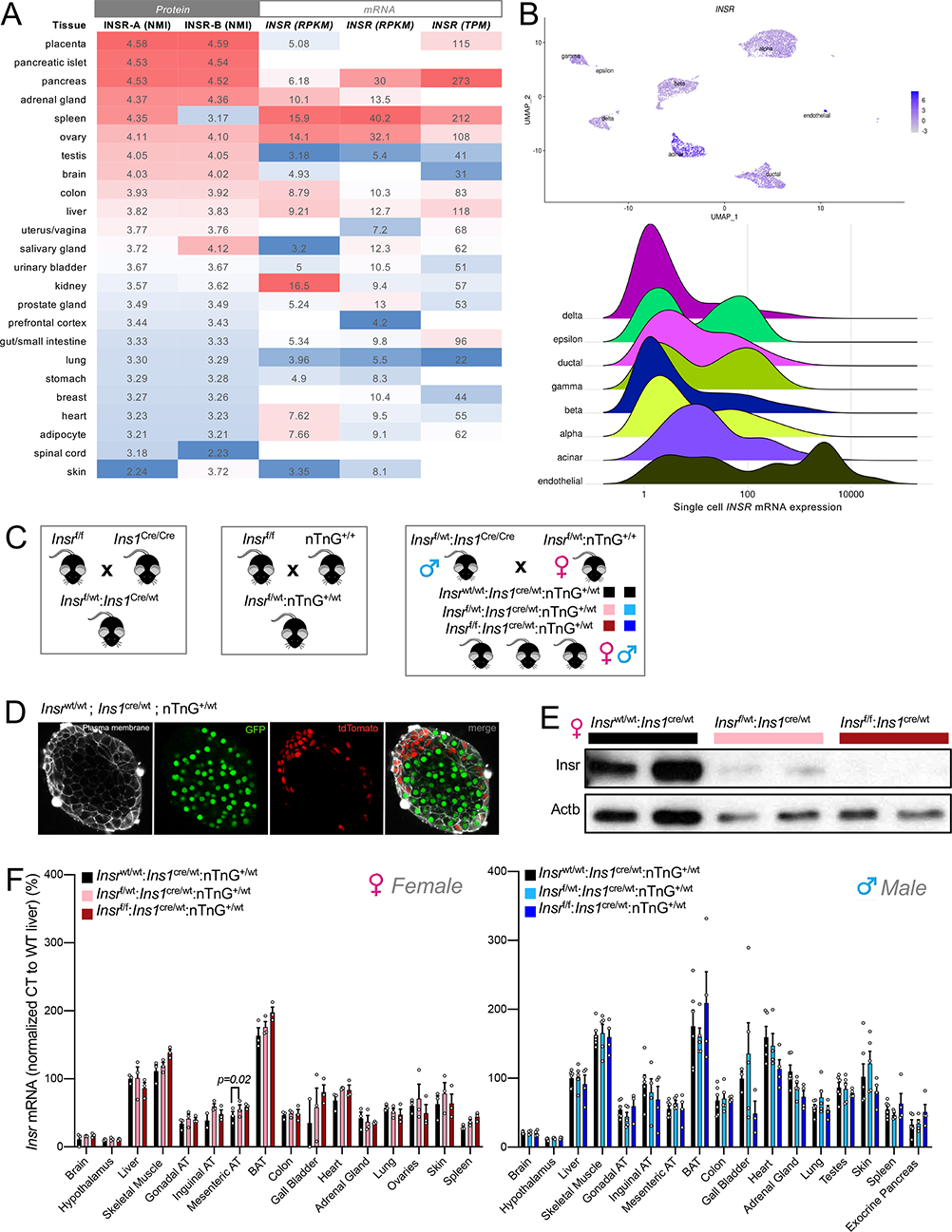
An animal model to examine the role of abundant Insr in β-cells. **(A)** INSR protein (isoform A and B) and *INSR* mRNA expression across human tissues. *Left 3 columns* show INSR isoform A and B protein abundance expressed as normalized median intensity (NMI; www.proteomicsdb.org). *Right 3 columns* show total *INSR* mRNA expression in the same tissues (where available) extracted from 3 databases. *Left* to *right*, mRNA data from human proteome atlas (HPA) and genotype tissue expression project (GTEX) are shown in reads per kilobase million (RPKM). Data from the FAMTOMS database is shown as transcripts per kilobase million (TMP). **(B)** Human insulin receptor expression in islet cell subtypes extracted from an integrated dataset of human single cell RNA seq data (see Methods). Normalized expression levels are shown in UMAP space (*top*) and as a ridge plot on a log scale (*bottom*). The height of the ridge indicates the frequency of cells at a given expression level. **(C)** Breeding strategy for generating β-cell specific β*Insr*^KO^, β*Insr*^HET^, and littermate control Cre-only mice. **(D)** Robust Cre recombination verified by imaging the nTnG reporter allele in an isolated islet from an *Ins1*^Cre/wt^;nTnG mouse. **(E)** Western blot of Insr protein in islets isolated from β*Insr*^KO^, β*Insr*^HET^, and littermate control Cre-only mice. **(F)** *Insr* mRNA expression across tissues in 16 week-old LFD-fed control, β*Insr*^HET^, and β*Insr*^KO^ mice assessed by qPCR (n=3 in females, n=5-6 in males).

### β-cell specific Insr deletion with Ins1^Cre^mice

We next sought to examine the function of the Insr, and the consequences of β-cell-specific insulin resistance/sensitivity, using an *in vivo* β-cell specific knockout mouse model. To limit recombination of the floxed *Insr* allele to pancreatic β-cells, we used a Cre allele knocked into the endogenous *Ins1* locus which, unlike *Ins2* promoters, drives specific expression in β-cells^30^. Experimental *Insr*^f/f^;*Ins1*^cre/wt^;nTnG (β*Insr*^KO^) and *Insr*^f/wt^;*Ins1*^cre/wt^;nTnG (β*Insr*^HET^) mice and littermate control *Insr*^wt/wt^;*Ins1*^cre/wt^;nTnG mice were generated using a breeding scheme to ensure consistency of the Cre donor parent (Fig. 1C). Cre-recombinase efficiency was assessed using the nuclear TdTomato-to-nuclear EGFP (nTnG) lineage trace reporter^32^ and found to be robust on the *Insr*^f/f^;*Ins1*^cre/wt^;nTnG genetic background (Fig 1D). We confirmed by Western blotting that *Insr* protein was almost completely absent from β*Insr*^KO^ islets and partially reduced from β*Insr*^HET^ (Fig. 1E). qPCR showed that *Insr* mRNA was not decreased in any of the 18 tissues examined, including the whole brain or hypothalamus specifically, hence we did not perform Western blots for other tissues (Fig. 1F). Together with other published data on *Ins1*^cre^ mice, these findings strongly suggest that *Insr* deletion with *Ins1*^cre^ is robust and sufficiently specific to pancreatic β-cells.

### Loss of β-cell Insr alters gene expression in purified β-cells

To establish a baseline gene expression profile of our β-cell specific *Insr* knockout model, we performed an unbiased analysis of gene expression in ∼100 FACS purified GFP-positive β-cells labelled with the nTnG reporter isolated from both female and male mice (Fig. 2A). We found that *Insr* mRNA expression is similar female and male β-cells (Fig. 2B). We confirmed a similar reduction in *Insr* in female and male β*Insr*^KO^ β-cells, with an intermediate phenotype in found in the β*Insr*^HET^ β-cells (Fig. 2B). We did not observe a compensatory change in *Igf1r* mRNA expression (Fig. 2C). After excluding samples with insufficient β-cell purity and analyzing across both sexes, RNA sequencing revealed significant differences in the expression of 12 genes between β*Insr*^KO^ β-cells and wildtype β-cells (Fig. 2C,D, S1). However, when we analyzed female and male β*Insr*^KO^ β-cells separately, we identified sex-specific gene expression changes (Fig. 2D-F). In female-only analysis, 5 genes (including 2 pseudo-genes) were differentially expressed, while in male-only analysis, 64 genes were differentially expressed with an adjusted *p* value of > 0.05. At this cut-off, there was no overlap between the sex-specific gene expression patterns. Although our study was not designed or powered for direct comparisons between sexes, these highlight the importance of considering both sexes separately.

**Figure 2.**
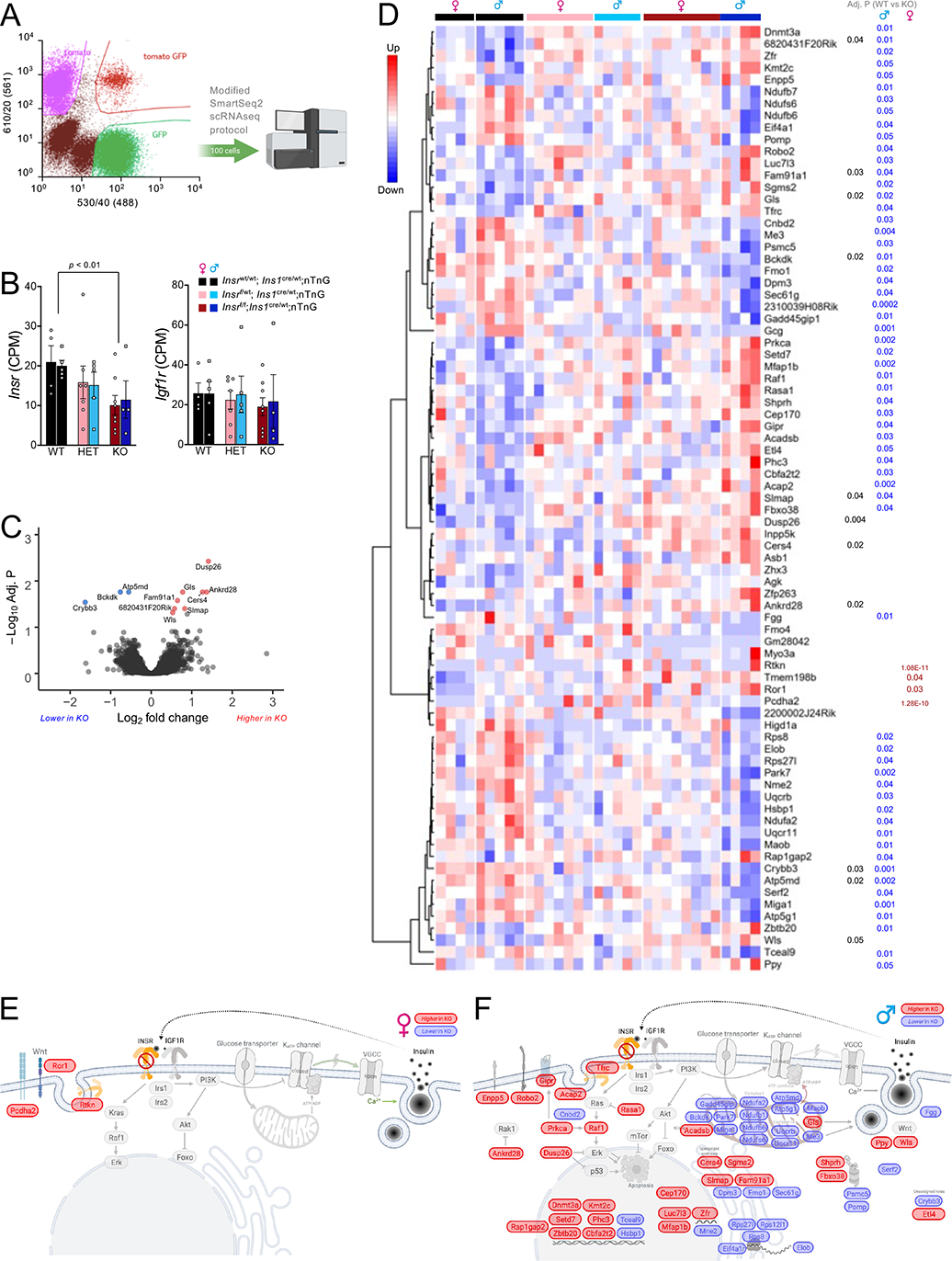
Transcriptomic analysis of purified *Insr* deficient β-cells. **(A)** Workflow for RNA sequencing of FACS purified GFP-positive β-cells (100 per group) from β*Insr*^KO^ and littermate control mice. **(B)** *Insr* and *Igf1r* mRNA levels. **(C)** Volcano plot and significantly differentially genes across both sexes. **(D)** Heatmap of all genes that were differentially expressed between any group, clustered by similarity. *P* values are shown on the right for the β*Insr*^WT^ to β*Insr*^KO^ comparison for all, males, and females. **(E)** Differentially expressed genes between female β*Insr*^WT^ and β*Insr*^KO^ cells overlaid on a diagram of their predicted functional roles and subcellular locations. **(F)** Differentially expressed genes between male β*Insr*^WT^ and β*Insr*^KO^ cells overlaid on a diagram of their predicted functional roles and subcellular locations.

### Loss of β-cell Insr increases β-cell excitability

Insulin administration has been reported to open β-cell KATP channels, mediating negative feedback on insulin secretion ^33^. Lack of this endogenous insulin action through Insr would therefore be expected to lead to β-cell hyper-excitability in our mouse model in the context of high glucose. Indeed, electrophysiological analysis of single β-cells from female β*Insr*^KO^ mice confirmed a significant increase in action potential firing frequency during glucose stimulation, when compared to control β-cells, with no differences in resting potential, firing threshold, or action potential height (Fig. 3A,B). The reversal potential was right-shifted in β*Insr*^KO^ β-cells, further suggesting reduced K^+^ conductance (Fig. 3C). Hyper-excitability was not observed in *Insr* knockout β-cells from male mice, at the age we studied (Fig. 3D). We did not observe a statistically significant difference in depolarization induced exocytosis in single cells from either sex (Fig. 3E,F), suggesting that the late stages of insulin granule exocytosis are not altered under these conditions.

**Figure 3.**
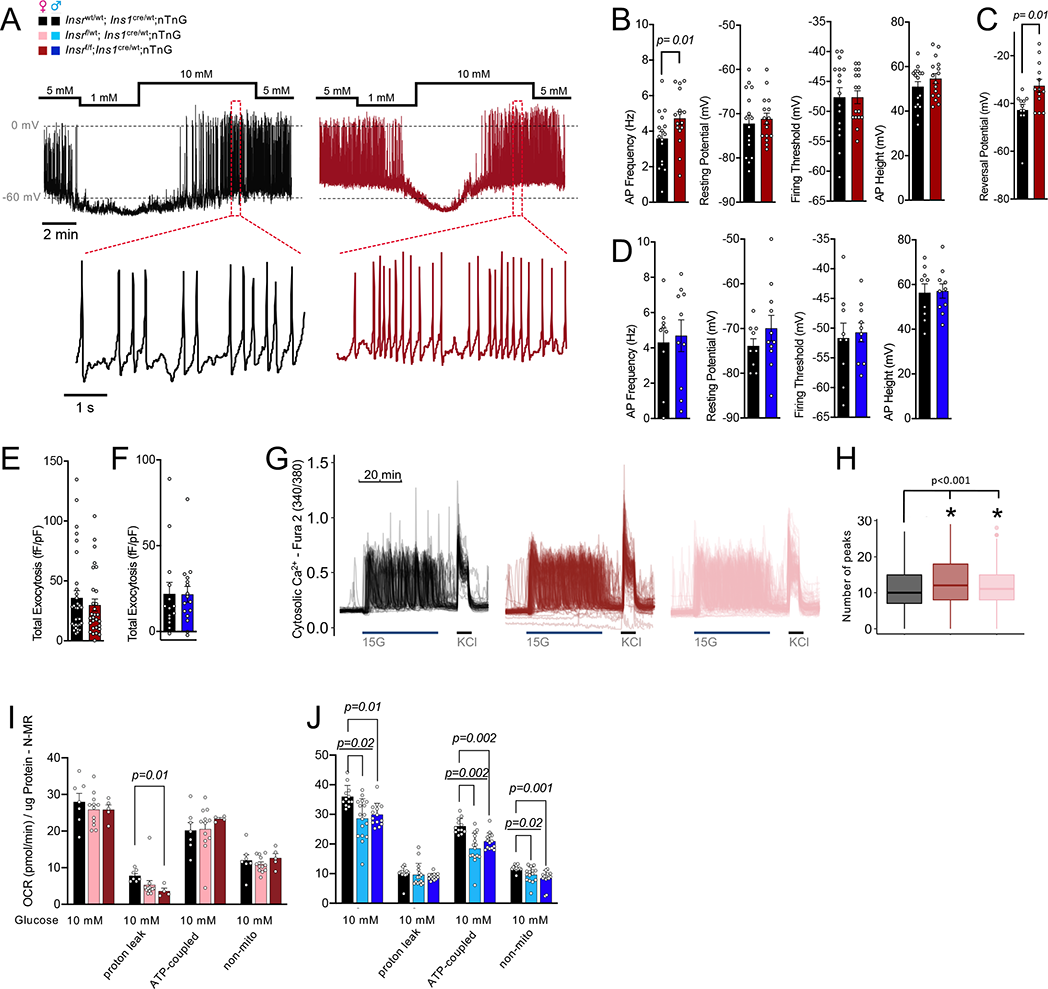
β-Cells lacking *Insr* have increased action potential and calcium oscillation frequencies. **(A)** Representative traces of action potential firing in β-cells from 16 week-old chow-fed mice. Glucose changed as indicated. **(B,C)** Quantification of action potential properties and reversal potential during the 10mM glucose phase in patch clamped dispersed β-cells. Data analyzed using un- paired t-test (n=16-17 cells, from 3 mice per group). (**D**) Quantification of electrophysiological properties in islets from male mice. **(E,F)** β-Cell exocytosis measured as increased membrane capacitance normalized to initial cell size (fF/pF) over a series of ten 500 ms membrane depolarizations from -70 to 0 mV (n= control 32 cells, 29 β*Insr*^KO^ cells, from 3 pairs of mice). **(G)** Ca^2+^ dynamics measured in dispersed islet cells (Fura-2 340/380 ratio) treated as indicated from a baseline of 3 mM glucose). **(H)** Quantification of glucose induced Ca^2+^ oscillation number (n=3523 cells). ANOVA with correction for multiple comparisons using Tukey’s method. Additional quantification of these traces can be found in Fig. S2A. **(I,J)** Oxygen consumption rate data of dispersed islets from 16 week-old chow-fed control, β*Insr*^HET^, and β*Insr*^KO^ mice. Data analyzed with mixed effects model from independent islet cultures from 3 male and 1 female mice.

Next, we analyzed Ca^2+^ responses to 15 mM glucose in thousands of Fura-2-loaded dispersed islet cells and analyzed the data with an adaptation of our TraceCluster algorithm^34^. In agreement with the electrophysiological data, *Insr* knockout β-cells from female mice exhibited a significantly greater number of oscillation peaks within the glucose stimulation period compared to control cells (Fig. 3G,H, S2A). A similar increase in excitability was observed in β*Insr*^HET^ β-cells. This was not associated with significant differences in the intensity or gross localization of Glut2 protein using immunofluorescence imaging (Fig. S2B).

We examined mitochondrial function using the Seahorse bioanalyzer in the context of 10 mM glucose. In dispersed islet cells isolated from female mice, there were no significant differences between genotypes in oxygen consumption rate, with the exception of a reduced proton leak (Fig. 3I). In contrast, dispersed islet cells from both β*Insr*^KO^ and β*Insr*^HET^ males had a significant reduction in glucose-stimulated oxygen consumption rate compared to controls. Oligomycin injection revealed that this included a decrease in ATP-linked respiration, suggesting reduced ATP production in islet cells from male mice lacking *Insr* (Fig. 3J). Notably, RNA sequencing showed that male, but not female, β*Insr*^KO^ had significantly decreased expression of many key mitochondria-related genes (e.g. *Bckdk*, *Miga1*, *Park7, Me3*), including 4 components of the NADH:ubiquinone oxidoreductase complex (complex 1 of the electron transport chain; *Ndufb7*, *Ndufs6*, *Ndufb6*, *Ndufa2*), 2 components of ubiquinol-cytochrome c reductase complex (electron transport chain complex 3; *Uqcr11*, *Uqcrb*), and 2 components of ATP synthase (*Atp5g1*, *Atp5md*) (Fig. 2D,F). Male-specific transcriptomic consequences of *Insr* loss could account for the sex difference in mitochondrial function that would be expected to impact ATP-dependent membrane potential and Ca^2+^ oscillations. Collectively, these experiments demonstrate that β-cells lacking *Insr* have increased electrical activity, so long as their mitochondrial ATP production is not impaired. Overall, our data support the concept that insulin normally has a negative feedback influence on excitability in the context of 10 mM glucose.

### Loss of β-cell Insr causes insulin hypersecretion in the context of stimulatory glucose

Insulin secretion is driven by electrical excitability, so we next carefully examined the effects of partial and full β-cell *Insr* deletion on secretory function employing multiple orthogonal *in vitro* and *in vivo* assays. We used islet perifusion to examine the dynamics of insulin secretion *ex vivo* at rest (3 mM glucose) and in response to 20 mM glucose or 10 mM glucose, as well as direct depolarization with 30mM KCl. Islets from female 16 week-old β*Insr*^KO^ and β*Insr*^HET^ mice secreted more insulin in response to 20 mM glucose and 30 mM KCl compared to islets from control mice (Fig. 4A). No significant differences were observed at low glucose (Fig. 4A). Consistent with our electrophysiology data, we did not observe differences in islets from males of the same age and on the same diet (Fig. 4B).

**Figure 4.**
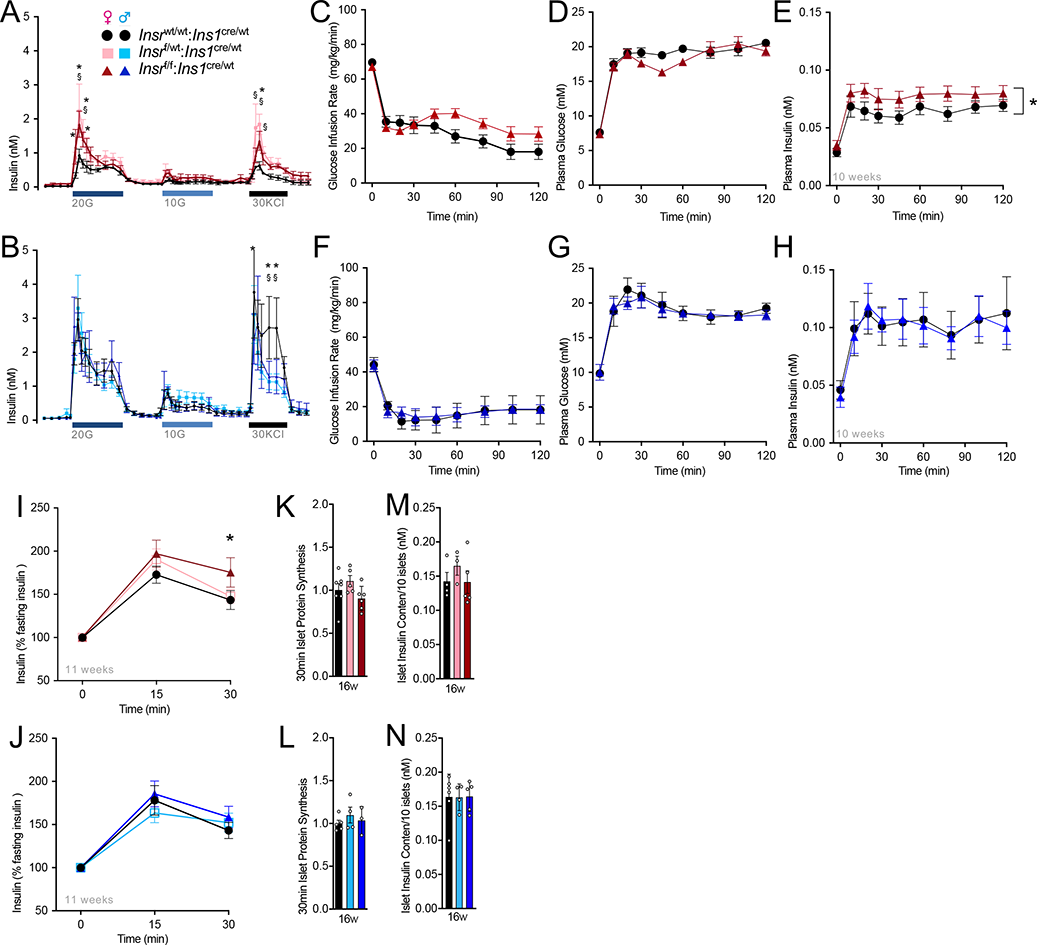
*Insr* knockout increases glucose-stimulated insulin secretion *in vitro* and *in vivo*. **(A,B)** Perifused islets isolated from 16 week old chow-fed control, β*Insr*^HET^, β*Insr*^KO^ female (n=3,3,5), and male (n=6,5,5) mice were exposed to 20 mM glucose (20G), 10 mM glucose (10G), and 30 mM KCL (KCL) from a baseline of 3 mM glucose. Data analyzed using repeated measures mixed effects model. Quantification of area under the curve (AUC) is shown for 1^st^ phase and 2^nd^ phase during 15 mM glucose stimulation, total response during 10 mM glucose stimulation, and during 30mM KCL stimulation. AUC’s were analyzed with 1-way ANOVA analysis. **(C-H)** Glucose infusion rates, plasma glucose levels, and plasma insulin levels during 2-h hyperglycemic clamps in awake LFD-fed, 10-week old control and β*Insr*^KO^ female (n=14, 15) and male (n=4, 4) mice. Data were analyzed using repeated measures mixed effects model. **(I,J)** Insulin levels (% basal insulin) following a single glucose injection (2g glucose/kg body mass,*i.p*) of 11 week old LFD-fed control, β*Insr*^HET^, β*Insr*^KO^ female (n=33,34,23) and male (n=22,29,17) mice. Data were analyzed using repeated measures mixed effects model. **(K,L)** 30min islet protein synthesis measured by S35 labeling in islets isolated from control, β*Insr*^HET^, β*Insr*^KO^ female (n=7,5,6) and male (n=5,4,3) mice. Data were analyzed by 1-way ANOVA. **(M,N)** Insulin content of 10 islets isolated from control, β*Insr*^HET^, β*Insr*^KO^ female (n=4,3,5) and male (n=5,4,5) mice. Data were analyzed by 1-way ANOVA.

This potentiation of high glucose-stimulated insulin secretion *ex vivo* in the complete and partial *Insr* knockout β-cells, led us to examine how insulin levels were affected by glucose stimulation *in vivo* using the hyperglycemic clamp technique in awake mice. For this cohort of mice, there were no significant differences in body mass (control 20.5 +/- 0.5g n=8 vs β*Insr*^KO^ 20.8 +/- 0.4g n=10), lean mass (control 17.5 +/- 0.4g vs β*Insr*^KO^ 17.3 +/- 0.2g), or fat mass (control 1.8 +/- 0.3g vs β*Insr*^KO^ 2.2 +/- 0.2g). Glucose infusion rates were adjusted in order to reach hyperglycemic levels (∼19 mM) in β*Insr*^KO^ and wild type control mice. Interestingly, slightly higher glucose infusion rates were necessary in female β*Insr*^KO^ mice in comparison to control mice in order to reach similar hyperglycemic levels (Fig. 4C). In accordance with our *ex vivo* insulin secretion data, glucose-stimulated insulin secretion was higher in female, but not in male β*Insr*^KO^ mice compared with control mice (Fig. 4C-H). We further tested whether *in vivo* insulin secretion would be potentiated after a single bolus of glucose in mice with reduced β-cell *Insr*. At 11 weeks of age, plasma insulin response, relative to baseline, was significantly elevated 30 min after i.p. injection of 2g glucose/kg body mass in female, but not in male, β*Insr*^KO^ mice compared to controls (Fig. 4I, J). In accordance with the electrophysiology, Ca^2+^ oscillation, and islet perifusion data, we detected no statistical difference between female genotypes in fasting plasma insulin *in vivo* at multiple ages (Fig. S3). Together, these experiments suggest that β-cell Insr can play suppressive role in glucose-stimulated insulin secretion without much impact on basal insulin secretion.

### Effects of β-cell Insr loss on insulin production, storage, processing and clearance

Insulin and insulin signaling can modulate protein synthesis in many cell types and we have previously provided evidence that soluble cellular insulin protein transiently increases in human islet cell cultures treated with exogenous insulin ^13^. To assess the quantitative contribution of Insr signaling to insulin production and mRNA translation rates we measured total islet insulin content after acid-ethanol extraction and examined total protein synthesis rate using S^35^- methoioine/cysteine pulse labelling in isolated islets. Total insulin content and protein synthesis in isolated islets were unaffected by *Insr* deletion under these basal glucose conditions (Fig. 4K-N).

To investigate the role of Insr signaling on β-cell stress and insulin clearance, we conducted analysis of plasma proinsulin to C-peptide ratios, and C-peptide to insulin ratios in the fasting state (4h) across multiple ages in both male and female mice, and on multiple diets (Fig. S4). While many of these parameters changed with age, no statistical differences between genotypes were seen in any of these parameters of mice fed either a low-fat diet (LFD) or a high fat diet (HFD). A trend towards lower insulin clearance was observed in LFD-fed female β*Insr*^KO^ mice in comparison to wild type control mice at 7 weeks. Collectively, these experiments show that β-cell insulin receptor signaling has only a small, if any, effect on insulin processing and clearance at baseline glucose conditions.

### Beta-cell area and hyperglycemia-induced proliferation in mice lacking β-cell Insr

Our RNA sequencing data revealed pathways downstream of *Insr* signaling that may affect β-cell proliferation capacity (e.g. *Raf1*)^16^, specifically in male mice (Fig. 2F). Thus, we examined baseline β- cell area and proliferation reserve capacity. Islet architecture and β-cell-to-α-cell ratio were not obviously perturbed (Fig. 5A). We did not detect significant differences associated with genotype in β-cell area in either female or male mice, at 13, 42 weeks (LFD) or 54 weeks (HFD) of age (Fig. 5B-E). In comparison to control mice, HFD fed female β*Insr*^KO^ mice had a tendency toward a smaller β-cell area at 54 weeks of age that were consistent with tendencies towards lower plasma insulin (Fig. 5D, S3C), proinsulin and C-peptide levels (Fig. S4C). These data suggest that Insr may help support age-dependent β-cell expansion under some conditions.

**Figure 5.**
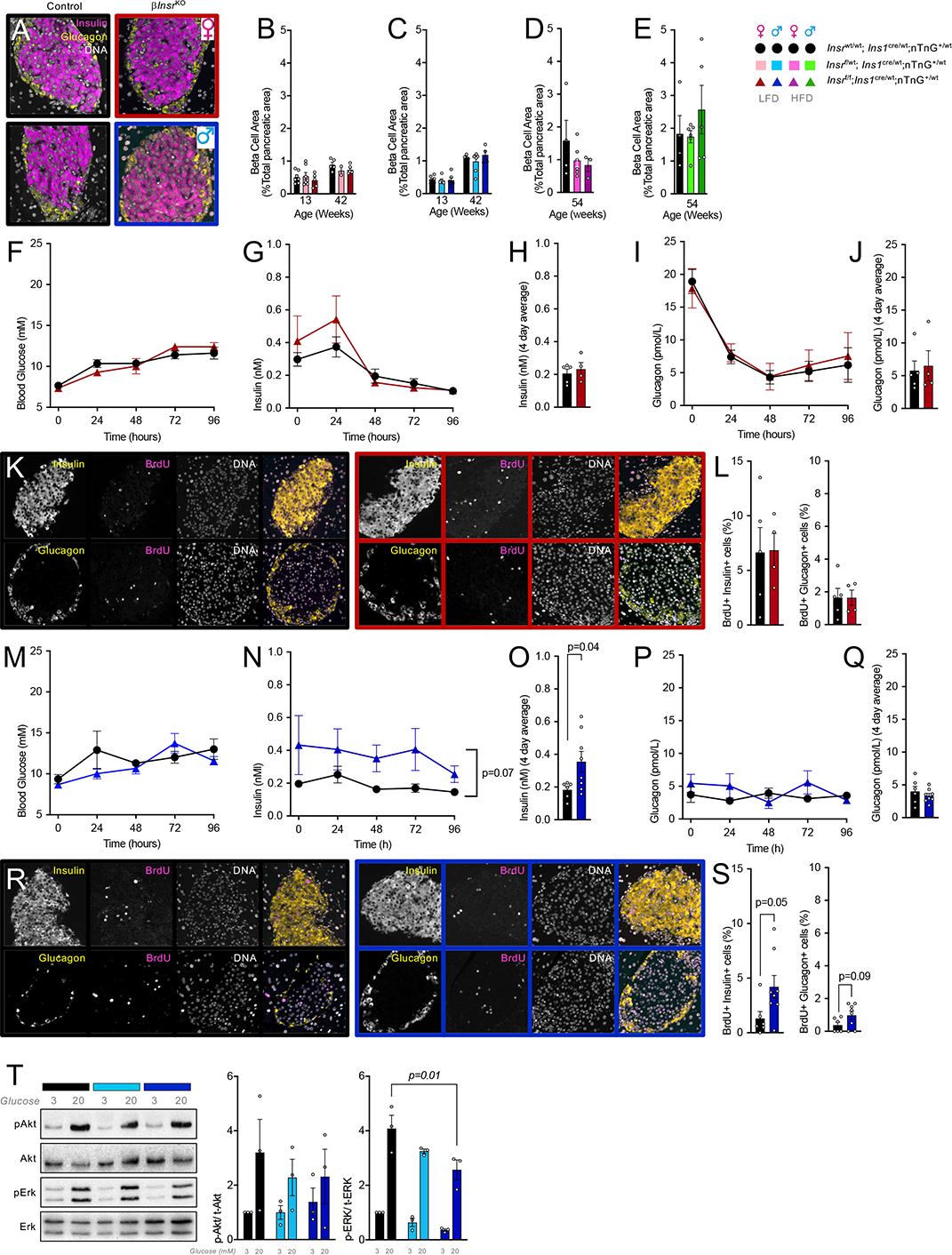
Islet cell proliferation and relative area in mice lacking β-cell *Insr*. **(A)** A representative image showing islet architecture via staining for insulin, glucagon and DNA. **(B-E)** β-Cell area shown as a percentage of total pancreatic area. Data were analyzed by 1-way ANOVA analyses. **(F-S)** 4-day *in vivo* glucose infusion in female (n=5 control, 4 β*Insr*^KO^ ) and male (n=6 control, 8 β*Insr*^KO^) mice. **(F,M)** Tail blood glucose and **(G,N)** insulin data were analyzed using repeated measures mixed effects model. **(H,O)** 4 day average plasma insulin data were analyzed by an unpaired t-test. **(I,P)** Plasma glucagon (tail blood) data were analyzed using repeated measures mixed effects model. **(J,Q)** 4 day average plasma insulin data were analyzed by an unpaired t-test. **(K,R)** Representative single channel and merge images showing islets stained for insulin, glucagon, BrdU and DNA from LFD-fed female and male controls and female β*Insr*^KO^ and male β*Insr*^KO^ mice following 4day glucose infusion. **(L,S)** Quantification of BrdU+ insulin+ cells and . BrdU+ glucagon+ cells. Data were analyzed by an unpaired t-tests. **(T)** Representative western blot image and quantification of islet lysate from male *Insr*^wt/wt^;*Ins1*^cre/wt^;nTnG (black bar, n=3), *Insr*^f/wt^;*Ins1*^cre/wt^;nTnG (light blue bar, n=3), *Insr*^f/f^;*Ins1*^cre/wt^;nTnG (dark blue bar, n=3) mice treated with 3 mM or 20 mM glucose. Data were analyzed by 1-way ANOVA.

Prolonged hyperglycemia can stimulate β-cell proliferation in adult mouse β-cells^35^, but whether this requires intact insulin receptor signaling remains controversial. To examine the role of *Insr*-mediated signaling on hyperglycemia-induced β-cell proliferation, we performed 4-day hyperglycemic infusions in β*Insr*^KO^ and wild type control mice. In female mice, hyperglycemia (>10 mM; Fig. 5F) resulted in mildly elevated insulin secretion in β*Insr*^KO^ relative to control mice for the initial 48 h, which was not sustained for the duration of the experiment (Fig. 5G,H), while glucagon levels declined similarly in both genotypes (Fig. 5I,J). There was no effect of *Insr* deletion on hyperglycemia-induced proliferation of either β-cells or α-cells in females (Fig. 5K,L). In male mice, 96 h of hyperglycemia resulted in sustained hyperinsulinemia in β*Insr*^KO^ mice (Fig. 5M-O), with no differences in circulating glucagon (Fig. 5P,Q). In male mice lacking β-cell *Insr*, this manipulation was associated with significantly more β-cell proliferation (Fig. 5R,S). The fact that we did not observe a suppression of glucose-induced proliferation of β-cells lacking *Insr* prompted us to determine the degree to which the, broadly defined, insulin signaling pathway was inhibited in our model. Indeed, glucose-induced Akt phosphorylation, shown by western blot of whole islet lysate, was statistically unaffected, and glucose-induced Erk phosphorylation was only reduced ∼50% in *Insr* knockout β-cells (Fig. 5T). It is likely that the *Igf1r* or another receptor tyrosine kinase or a receptor tyrosine-kinase-independent mechanism initiates parts of intracellular post-receptor ‘insulin signaling’ in the absence of *Insr*. Testing this hypothesis in the future will require truly β-cell specific double deletion of *Insr* and *Igf1r*, and/or additional receptors.

### Modelling contributions of peripheral and β-cell specific insulin sensitivity to glucose homeostasis

The continuum between insulin sensitivity and resistance impacts multiple tissues, including pancreatic β-cells. We observed that β-cell-specific insulin resistance resulted in insulin hypersecretion in the context of unchanged β-cell mass. We next used mathematical modelling to generate quantitative predictions of the dependence of glucose tolerance on both β-cell and whole-body insulin resistance, both independently and synchronously. As described in the methods section, we modified the Topp model ^36^, adding insulin receptor-mediated negative feedback on insulin secretion, as indicated by our experimental data, with Sβ serving as the β-cell Insr-specific insulin sensitivity parameter (see equations in Fig. 6A-C; for β*Insr*^KO^ mice, Sβ = 0). Peripheral insulin sensitivity is represented by SP, (SI in the original Topp model). We used our hyperglycemic clamp data (Figs. 4C-E, S5A,B) to estimate Sβ in both female and male control mice and found Sβ,female to be significantly different from zero (Sβ,female = 3.4 +/- 1.5 nM-^1^, p = 10^-25^) and significantly different from Sβ,male (p = 10^-24^). Sβ,male was not significantly different from zero (Sβ,male = -0.05 +/- 1.0 nM^-1^, p = 0.7). *In silico* glucose tolerance tests found that decreased β-cell insulin-sensitivity (Sβ) (similar to β*Insr*^KO^ mice) corresponded with elevated peak and plateau insulin secretion (Fig. 6D). As expected, the computations predicted more rapid clearance of blood glucose (Fig 6E). Analysis of areas under the curve for glucose and insulin resulting from *in silico* glucose tolerance tests while varying SP and Sβ indicated that β-cell insulin resistance should have a marked effect on insulin secretion and glucose tolerance, most dramatically in conditions of low peripheral insulin sensitivity (Fig. 6F,G). Next, we compared the predictions of this model with experimental results. We used the *in silico* AUCGlucose values as a function of both SP and Sβ combined with averaged experimental AUGC values to estimate SP as a function of age for the low-fat diet conditions (Fig. 6H,I). We found that male values of SP for wildtype and mutant were indistinguishable from each other while females showed significant differences from each other and from the male values at all ages. As expected, HFD led to reduced peripheral insulin sensitivity (Fig. 6J). Collectively, these simulations show how β-cell insulin sensitivity and peripheral insulin sensitivity may combine to the regulation of glucose tolerance.

**Figure 6.**
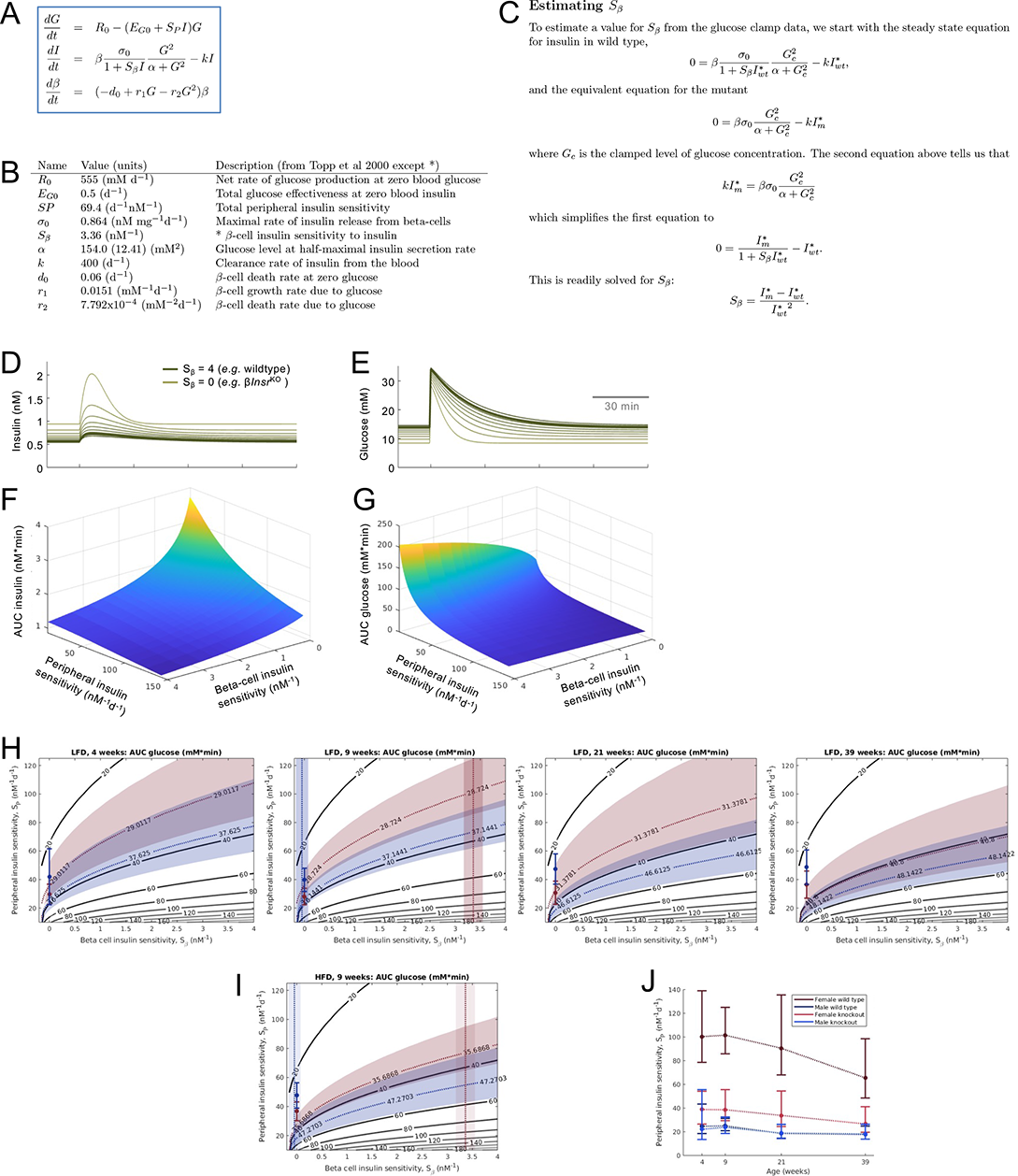
Mathematical modelling of insulin secretion and glucose tolerance. **(A,B)** Modified version of the Topp model. **(C)** Estimation of *Sβ* from hyperglycemic clamp data. **(D, E)** Simulations of the effects of reduced β-cell insulin sensitivity on glucose stimulated insulin release and glucose tolerance. **(F, G)** Relationship between the contributions of peripheral insulin sensitivity and β-cell insulin sensitivity to the glucose AUC and insulin AUC in the *in silico* glucose tolerance tests. **(H)** Modelled effects of peripheral insulin sensitivity and beta-cell insulin sensitivity on glucose AUC at different ages (topographical lines). Female response is shown in the *pink* shadow and male response in the *blue* shadow. The experimental effects of the knockout are shown as the single data point with SEM bars. (**I)** Effects of HFD at 9 weeks, where comparable glucose bolus was given. **(J**) Model-assessed change in peripheral insulin sensitivity with age, using experimental data.

### Context-dependent improvement in glucose tolerance with reduced β-cell Insr signaling

To test our theoretical model experimentally, we examined glucose tolerance in female and male

β*Insr*^KO^, β*Insr*^HET^, and control littermates at multiple ages between 4 and 52 weeks in the context of two diets. Significant improvements in glucose tolerance were observed in female mice with reduced Insr signaling at multiple ages, and in young males (Fig. 7). Consistent with our mathematical modelling that suggested a diminished contribution of β-cell insulin resistance to glucose homeostasis in the context of greater whole-body insulin resistance, we did not observe significant effects of genotype in older male mice, or mice of either sex fed an insulin-resistance-inducing HFD, in contrast to models with both β- cell and brain *Insr* knockout^20–22^. Thus, *Insr* deletion specifically in β cells had less effect on glucose tolerance in mice with already impaired pan-tissue insulin resistance, which we and others have shown increases with age and is more pronounced in male mice (Fig. 6J, S6).

**Figure 7.**
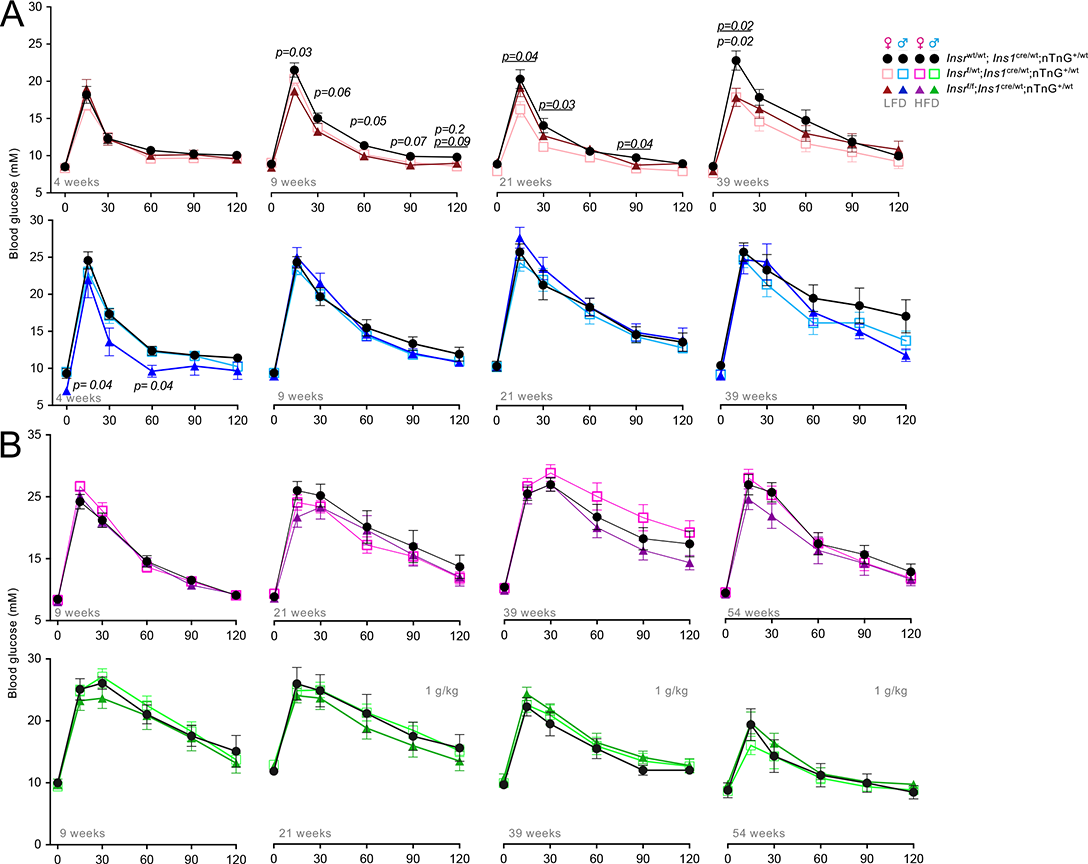
Glucose tolerance at multiple ages in β-cell specific *Insr* knockout mice fed 2 different diets. **(A)** Glucose tolerance tests of control, β*Insr*^HET^, β*Insr*^KO^ LFD-fed females (n4week=16,16,14; n9week=23,25,17; n21week=12,15,12; n39week=8,12,10) and males (n4week=10,18,5; n9week=17,30,11; n21week=8,14,8; n39week=9,7,13) **(B)** Glucose tolerance tests of control, β*Insr*^HET^, β*Insr*^KO^ HFD-fed female (n9week=17,17,14; n21week=12,14,12 n39week=14,16,10; n54week=10,13,11) and male (n9week=7,16,9; n21week=7,16,8; n39week=8,14,8; n54week=6,11,7) mice. All mice received a glucose bolus of 2g glucose/kg body mass *(i.p)* except older HFD-fed males, which received only 2g glucose/kg body mass *(i.p*). \**p- values are italicized* when β*Insr*^KO^ was compared to controls, p-values are underlined when β*Insr*^HET^ was compared to controls.

The multiple analyses conducted on both sexes, at different ages, and one two diets can be appropriately analyzed using Bayesian methods. Similar to the frequentist statistical analysis, we observed evidence for improvements in glucose tolerance in female mice with reduced Insr signaling at 9 and 39 weeks of age, and in males at 4 and 39 weeks of age (Fig. 8). In the context of high fat diet-induced insulin resistance, we only observed evidence for improved glucose tolerance in older female mice without β cell Insr signalling. Generally, this statistical analysis validates and extends the more commonly used statistical methods above.

**Figure 8.**
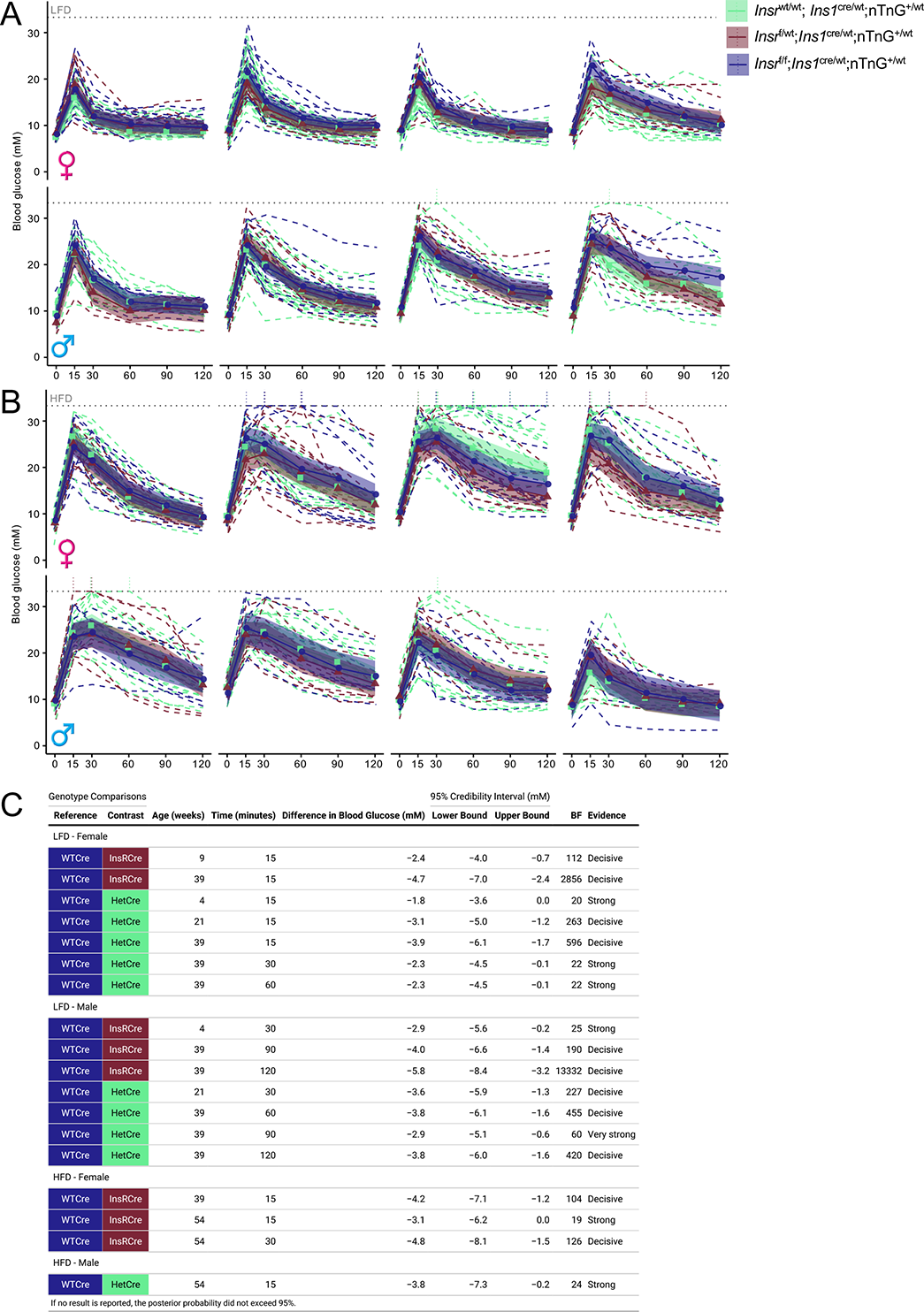
Bayesian regression modelling of glucose tolerance at multiple ages in β-cell specific *Insr* knockout mice fed 2 different diets. Same data as Figure 6. Glucose tolerance tests of control, β*Insr*^HET^, β*Insr*^KO^ LFD-fed female (n4week=16,16,14; n9week=23,25,17; n21week=12,15,12; n39week=8,12,10) and male (n4week=10,18,5; n9week=17,30,11; n21week=8,14,8; n39week=9,7,13) as well as HFD-fed female (n9week=17,17,14; n21week=12,14,12 n39week=14,16,10; n54week=10,13,11) and male (n9week=7,16,9; n21week=7,16,8; n39week=8,14,8; n54week=6,11,7) mice. All mice received a glucose bolus of 2 g glucose/kg body mass *(i.p)* except older HFD-fed males, which received only 1 g glucose/kg body mass *(i.p*). Data were analysed using Bayesian multilevel regression modelling. Dashed lines indicate individual mice, solid lines and points indicate estimates, and shading indicates 95% credible intervals. Dotted line at 33.3 mM indicates upper limit of detection of glucometer and vertical dotted lines rising above this line indicate where measurements were above the limit of detection for an individual mouse.

### Acutely improved glucose tolerance with inducible β-cell-specific Insr loss

The *Ins1*^cre^ allele results in pre-natal gene deletion^30^. To assess the effects of *Insr* deletion in adult β- cells, and to determine the role of *Insr* on a different genetic background and under different housing conditions, we also phenotyped multiple cohorts of male mice in which the *Insr*^f/f^ allele was recombined by the *Ins1* promoter-driven CreERT transgenic allele (commonly known as MIP-Cre) after injection with tamoxifen. In agreement with our observations in mice with constitutive loss of β-cell *Insr*, we found that glucose tolerance was significantly improved 4 weeks after β-cell-specific *Insr* deletion in male mice (Fig. 9). These differences were not maintained as the mice aged and became more insulin resistant. In these mice, there were no significant differences observed in fasting glucose (control 4.8 +/- 0.3mM n=7 vs β*Insr*^KO^ 4.4 +/- 0.2mM n=10), β-cell mass (control 1.4 +/- 0.3% n=3 vs β*Insr*^KO^ 2.1 +/- 0.2% n=5), or body mass (control 26.1 +/- 1.1g n=7 vs β*Insr*^KO^ 23.1 +/- 0.7g n=17. Collectively, these observations using an independent model and independent housing conditions lend support to our conclusion that the initial consequence of β-cell specific *Insr* deletion is improved glucose tolerance. This experiment also demonstrates that the role of Insr in β-cell function is not formally sex specific, just sex biased.

**Figure 9.**
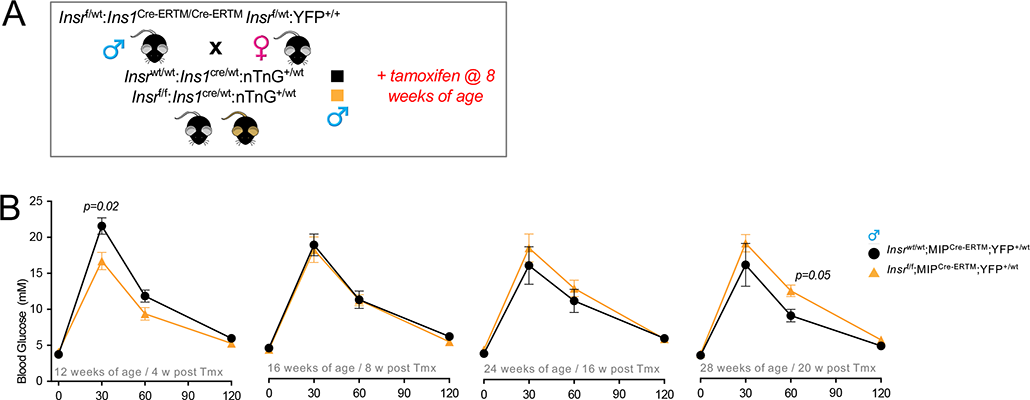
Glucose tolerance at in inducible β-cell specific *Insr* knockout mice. **(A)** Breeding and experimental designed for inducible β-cell specific Insr knockout mice. **(B).** Glucose tolerance of Chow- fed *Insr^wt^*^/wt^;MIP^Cre-ERTM^;YFP^+/wt^ (n=7) and *Insr*^f/f^;MIP^Cre-ERTM^;YFP^+/wt^ (n=17) mice were examined at 4, 8, 16 and 20 weeks after tamoxifen injection at 8 weeks of age (3 x 200 mg/kg tamoxifen over a 1-week period). Data were analysed using repeated measures mixed effects models.

### Peripheral effects of β-cell specific Insr loss

We and others have shown that even modest differences in hyperinsulinemia can have profound consequences for insulin sensitivity, adiposity, fatty liver, longevity and cancer^14, 37, 38^. Thus, we asked how the context-dependent glucose-stimulated insulin hyper-secretion induced by targeted β-cell specific insulin resistance may affect insulin sensitivity, adiposity, and body mass over time. Insulin sensitivity was assessed at multiple ages in the same mice. Interestingly, insulin sensitivity was significantly improved in 10-week-old female β*Insr*^HET^ mice compared to littermate controls without *Insr* deletion (Fig. S6A). On a high fat diet, male β*Insr*^KO^ and β*Insr*^HET^ mice had significantly improved insulin sensitivity compared to controls at 22 weeks of age (Fig. S6D). Longitudinal tracking of 4-h fasting glucose identified relative hypoglycemia in young LFD female β*Insr*^KO^ and β*Insr*^HET^ mice, older LDF male β*Insr*^KO^ and β*Insr*^HET^ mice, and across the tested ages in HDF female mice (Fig. 10A-D). Longitudinal tracking of body weight revealed that female mice with reduced β-cell Insr consistently weighed more than controls when fed a HFD (Fig. 10E-H), consistent with the known role of hyperinsulinemia in diet-induced obesity^14, 39^. We also examined the mass of several tissues at 13 weeks of age. Interestingly, liver mass was lower in both female and male mice lacking β-cell *Insr* (Fig. 10I,J). Pilot experiments suggested that liver Insr protein abundance may have been reduced in mice with partially or completely reduced β-cell Insr, in the context of the LFD but not the HFD (Fig. 10K). These data are consistent with the concept that modest hyperinsulinemia can drive down Insr levels and the concept that insulin signaling is a trophic signal for liver. Together, these data demonstrate that specifically preventing autocrine insulin feedback can have systemic effects on insulin sensitivity, body mass, and the size of some tissues. These changes may affect the eventual susceptibility to type 2 diabetes.

**Figure 10.**
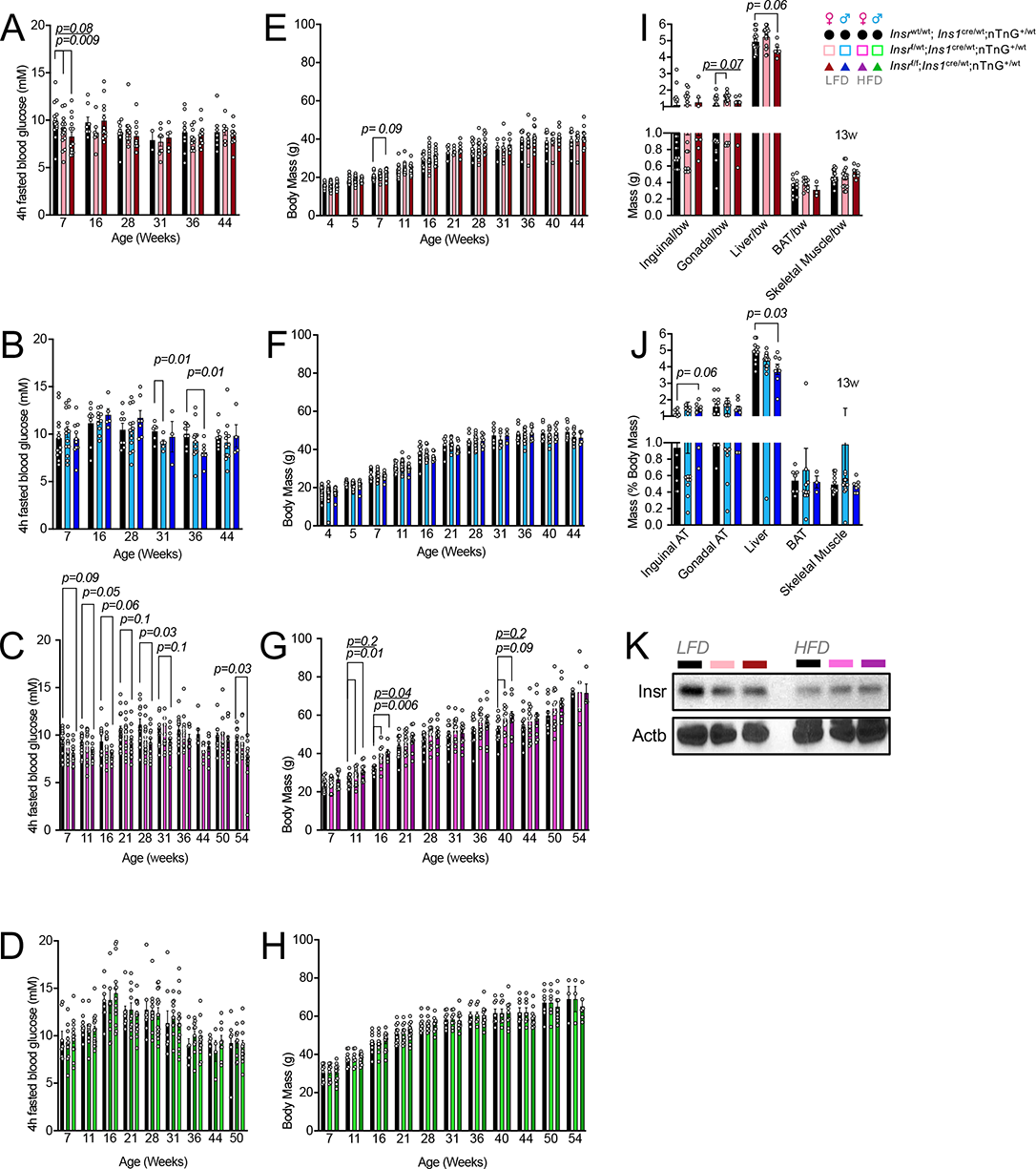
Effects of β-cell specific *Insr* deletion fasting glucose, body weight and organ weight. **(A-D)** Plasma glucose concentration after a 4-h fast in control, β*Insr*^HET^, and β*Insr*^KO^ mice at multiple ages in both LFD and HFD. **(E-H)** Longitudinally tracked body weight in control, β*Insr*^HET^, and β*Insr*^KO^ mice at multiple ages in both LFD and HFD. **(I,J)** Weights, as a percentage of the whole body, of inguinal adipose tissue, gonadal adipose tissue, liver, brown adipose tissue and skeletal muscle. Data were analysed using repeated measures mixed effects models. **(K)** Representative immunoblot of Insr (n = 4).

## Discussion

The goal of the present study was to establish the role of β-cell Insr on glucose homeostasis using specific genetic loss-of-function tools. We found that *in vivo Insr* gene deletion potentiated glucose-stimulated insulin secretion by increasing action potential firing and Ca^2+^ oscillation frequency, leading to improved glucose tolerance in insulin sensitive animals. Our data therefore suggest a model in which insulin inhibits its own secretion in a context-dependent manner and that this local negative feedback loop has physiological consequences for glucose tolerance.

Autocrine signaling in endocrine cells is generally a negative feedback^40^, with a few exceptions in specific conditions^41^. Given the abundance of Insr protein in islets and the physiological modulation of both insulin and Insr signaling in health and disease, autocrine and paracrine insulin signaling have been topics of interest and controversy for decades^6, 42^. While some have questioned whether local insulin levels are sufficient for signaling within the islet, mathematical modelling of insulin hexamer dissolution estimated that monomeric insulin within islets is in the picomolar range^43^, similar to the dose that maximally activates β-cell insulin signaling pathways^16, 44^. Consistent with a narrow range of responsiveness, our results also show that the contribution of autocrine insulin feedback to glucose homeostasis depends on whether mice are on a diet or at an age where insulin resistance is high, and potentially saturated in β-cells. Background genetics, diet, housing conditions, microbiome, or glucose concentrations could contribute to differences in observed phenotypes between our β-cell specific *Insr* knockout models and the frank diabetes reported for transgenic models that use fragments of the *Ins2* or *Pdx1* promoters to drive Cre-mediated β-cell *Insr* deletion^20–22^. However, we believe that the discrepancy is more likely due to depletion of *Insr* in key neuronal populations since both of *Ins2* and *Pdx1* are expressed in brain regions that influence glucose homeostasis^23, 45^. For example, *Insr* deletion in the brain with Nestin Cre causes insulin resistance^46^ and impairs the sympathoadrenal response to hypoglycemia^47^, while *Insr* knockout using AgRP Cre results in abnormal suppression of hepatic glucose production^48^. Our β*Insr*^KO^ mouse line is the most tissue specific model used to date for the study of autocrine insulin signaling and β-cell insulin resistance.

Using genetic and genomic tools, our work complements previous studies in humans and animal models. *Ex vivo* studies of perfused canine pancreata found an inhibitory autocrine effect of insulin ^49^. Similarly, exogenous insulin perfusion of canine pancreas *in situ* was shown to lead to reduced endogenous insulin production^50^. *In vivo* insulin infusion rapidly suppressed C-peptide levels in healthy men, but not those with obesity and presumably global insulin resistance^8^. In isolated human islets, perifusion studies showed that treatment with physiological doses of insulin had no effect on C-peptide release^51^, while static incubation experiments found only moderate potentiation of glucose-stimulated insulin secretion with super-physiological levels of insulin ^42^. Our conclusions are in line with the recent work of Paschen et al reporting that a high fat, high sucrose diet induced tissue-selective insulin resistance in β-cells, as well as profound hyperinsulinemia and β-cell hyper-excitability^5^. Our mechanistic finding that *Insr* knockout β-cells have increased action potential firing frequency is consistent with previous observations that insulin directly increased KATP currents via PI3-kinase signaling^33^ and that PI3-kinase inhibition with wortmannin potentiates glucose stimulated insulin secretion in normal, but not T2D, human islets^52^. In the islets from 16-week-old male *Insr* knockout mice, it seems likely that the significant reduction in ATP-production and differences in gene expression undercut this effect. Collectively, our studies further illustrate the molecular mechanisms of negative autocrine feedback in β-cells during high glucose stimulation.

Beyond the potentiation of glucose-stimulated insulin secretion, transcriptomic analysis of fully and partially Insr-deficient β-cells revealed a re-wiring of signaling pathways that could change how these cells respond to stresses (Fig. S1). For example, increased expression of the dual-specificity phosphatase *Dusp26* β*Insr*^KO^ cells is expected to modulate Erk signaling downstream of Insr. Dusp26 has been implicated as a negative regulator of β-cell identity and survival^53^. Increased *Ankrd28*, a regulatory subunit of protein phosphatase 6^54^, is expected to modulate inflammatory signaling. Increased ceramide synthase 4 (*Cers4*) would affect sphingolipid signaling and has shown to mediate glucolipotoxicity in INS-1 β-cells^55^. Increased mitochondrial glutaminase (*Gls*) would affect local glutamate synthesis, potentially affecting insulin secretion^56^. Increased Wls would support Wnt protein secretion. On the other hand, decreased branched-chain alpha-ketoacid dehydrogenase (*Bckdk*) would be expected to lower cellular levels of branched-chain amino acids. The consequences of decreased *Crybb3* are not clear in this context.

Our transcriptomic analysis pointed to robust sex dependent consequences of β-cell specific *Insr* deletion. Specifically in female β*Insr*^KO^ cells, we found increased expression of *Ror1*, a receptor tyrosine kinase implicated as a scaffold protein for caveolae-dependent endocytosis^57^, which mediates *Insr* internalization and survival in β cells^31^, an inhibitory scaffold of Rho GTPases (*Rtkn*), and the calcium-dependent adhesion protein (*Pcdha2*). The transcriptomic changes in male β*Insr*^KO^ cells were broad-based and included down-regulation of key genes controlling mitochondrial metabolism. There was also decreased *Hsbp1*, which links insulin signaling to longevity in *C elegans*^58^. The transferrin receptor (*Tfrc*) has been linked to insulin sensitivity and type 2 diabetes^59–61^. Male β*Insr*^KO^ cells also had increased *Rasa1*, which is a suppressor of Ras. Male β*Insr*^KO^ cells exhibited increased expression of signalling genes with known roles in β-cell function. Using MIN6 cells and primary β-cells from male mice, we previously showed that *Raf1* mediates many of insulin’s effects on β-cell survival, proliferation and β- cell function^16, 44, 62, 63^. Protein kinase C alpha (*Prkca*) is a direct upstream regulator of *Raf1* and may also stimulate the β-cell expression of incretin receptors such as *Gipr*^64^. *Robo2* plays important roles in β-cell survival ^65^ and islet architecture^66^. Male β*Insr*^KO^ cells also had up-regulation of pancreatic polypeptide (*Ppy*), but whether this reflects a change in β-cell differentiation status or the purity of sorted cells remains is unclear. Additional studies, beyond the scope of this investigation, will be required to assess the contribution of these differentially expressed genes towards the phenotype of female and male β*Insr*^KO^ mice.

The conditions under which these gene expression changes could manifest in long-term effects on β-cell proliferation or increased β-cell survival remain incompletely explored. Otani et al reported modestly reduced β-cell mass in non-diabetic *Ins2*-Cre transgenic Insr knockouts, which was exacerbated by diabetes, but they did not employ Cre controls^22^. Okada et al reported impaired compensatory β-cell proliferation in the context of high fat diet or liver *Insr* knockout in the same experimental groups^21^. These previous studies comparing β-cell knockout mice of Insr versus Igf1r, suggested a more important role for the former in β-cell proliferation and survival^21^. *Insr* over-expression experiments also support the idea that β-cells are key insulin targets^67^. In the present study, we were unable to identify conditions which would result in statistically significant differences in β-cell mass in mice with Insr deficiency, but variability was high and relatively few animals were studied at older ages. We have previously shown that the increase in β-cell mass resulting from high fat diet requires local insulin production and is independent of hyperglycemia^14^, consistent with the known direct anti-apoptotic and *Raf1*-dependent mitogenic effects of insulin *in vitro*^15, 16, 68^. These findings can be reconciled by proposing that high insulin concentrations within the islet are sufficient to activate remaining Igf1 receptors linked to *Raf1*-biased post-receptor signaling. We observed that sustained hyperglycemia and hyperinsulinemia over 4 days was associated with increased proliferation, but whether these are effects are mediated through Igf1r function compensation will require double receptor knockout experiments. The recent identification of a negative modulator of both Insr/Igf1r action in β-cells with profound effects on glucose homeostasis, supports the concept that double Insr/Igf1r inhibition inhibits β-cell proliferation^69^. It is also possible that hyperglycemia itself is a major driver of β-cell proliferation under these conditions, through Irs2-Creb signaling that may bypass Insr/Igf1r^18^. It has also been demonstrated that ∼80% of the gene expression changes attributed to glucose in MIN6 cells require full insulin receptor expression^17^. We have previously found that glucose cannot stimulate primary mouse β-cell proliferation when autocrine insulin signaling is blocked by somatostatin and that ‘glucose-induced’ Erk phosphorylation requires full insulin secretion^16, 44^. On the other hand, inhibiting Insr in mouse islets with S961 or shRNA did not block glucose-induced proliferation of cultured β-cells ^19^. Additional future studies will be required to resolve this controversy.

Early hyperinsulinemia is a feature of β-cell *Insr* knockout models on multiple genetic backgrounds^20–22^, including the present study. Loss of Irs1 or Akt function results in basal hyperinsulinemia and, in some cases, increased β-cell mass^70, 71^, mimicking the early stages in human diabetes. Our experiments begin to shed light on the systemic consequences of the hyperinsulinemia caused by β-cell-specific insulin resistance, which may be an early event in the pathogenesis of type 2 diabetes. Human data suggests that β-cell insulin resistance can be found in the obese state prior to hyperglycemia^8^. We and others have shown that hyperinsulinemia contributes to insulin resistance and obesity^14, 39^, through multiple mechanisms including the down-regulation of insulin receptors^72^. We observed propensity for excessive diet-induced body mass gain in mice with β-cell specific insulin resistance, as well as Insr protein down-regulation in the liver. Thus, β-cell defects such as impaired autocrine feedback through Insr may contribute to insulin hypersecretion and accelerate the early stages of type 2 diabetes. In the later stages, the lack of pro-survival insulin signaling, perhaps in combination with other molecular defects, may contribute to failures in β-cell compensation and survival, thereby further accelerating the course of the disease^17, 20, 21, 73, 74^.

Our studies illustrate the power of using both females and males to study integrated physiology, although it was not designed to test specifically for sex differences. Indeed, β-cell *Insr* loss led to increased β-cell action potentials, calcium oscillations, and glucose-stimulated insulin secretion in 16- week-old female mice, but not in age-matched males. Although significant sex differences have been reported at the transcript level for many genes in mouse and human islets^75–78^, no intrinsic sex differences of *Insr* mRNA levels were reported in sorted β-cells^77^, and we confirmed this in our analysis of sorted β-cells (Fig. 2B). Given that we found identical *Insr* reduction in both sexes of β*Insr*^KO^ mice, the sex differences in glucose-stimulated action potential firing rate, Ca oscillations, and insulin secretion must come from the sex-specific downstream transcriptomic consequences we identified (Fig. 2D-F). Of particular note, we identified a group of genes critical for mitochondrial ATP generation that were only significantly reduced in male β-cells, consistent with reduced ATP-coupled oxygen consumption at 10 mM glucose we measured in male, but not female, islets (Fig. 3K). This is likely to account for a least some of the observed sex difference. It is also possible that control male, but not female, β-cells were already maximally insulin resistant when studied. Given the abundance of data showing more pronounced insulin resistance in males ^79, 80^, this could be another reason for female-specific response to β-cell *Insr* knockout. To this point, glucose tolerance is improved in 4-week-old β*Insr*^KO^ males, an age at which control males remain insulin-sensitive, and in male mice with acute loss of *Insr*. More work will be needed to confirm this possibility, and to determine factors in addition to sex hormones and sex chromosomes that impact these sex differences in insulin sensitivity and glucose homeostasis. The sex-specific nature of phenotypes arising from our genetic manipulation of *Insr* in β-cells highlights the importance of including both sexes to accurately interpret data and to draw conclusions that will apply to both sexes. Additional research will be required to fully elucidate the molecular mechanisms underlying the myriad of sex differences in β-cell physiology.

While our study is comprehensive and employs the best genetic tools available today, this work has limitations. *Ins1*^Cre^ is the most β cell-specific Cre deletion strain available today ^30^, but this fact does not preclude off-tissue effects that have yet to be discovered. Cre recombinase itself is not totally benign. These facts and our detailed comparison of *Ins1*^wt/wt^ mice with *Ins1*^Cre/wt^ mice (Figs. S9-12), highlight the importance of the Cre control group we employed throughout our studies. Recently, it was reported that some colonies of *Ins1*^cre^ mice exhibited some silencing via DNA hypermethylation at the *Ins1* locus and this was suggested as an explanation for discordance between the phenotypes compared to gene deletions using Pdx1-Cre and Ins2-Cre transgenic lines ^81^. In our study, we observed virtually complete recombination in β-cells and no evidence for off-tissue *Insr* deletion. We believe a major source of discrepancy with previously reported phenotypes stems from the propensity of previous promoter transgenic strains to recombine in the brain and robust expression of *Insr* throughout the brain. Another caveat of our experiments using β*Insr*^KO^ mice is that *Insr* is expected to be deleted from β-cells starting in late fetal development^30^. In our hands, tamoxifen-inducible *Ins1*^CreERT^ mice have insufficient β-cell specific recombination for *in vivo* physiological studies. Our validation experiments using the tamoxifen-inducible *Ins1*-CreERT address this limitation and confirm that β-cell insulin resistance improves glucose tolerance, at least under the initial insulin sensitive conditions. It should also be noted that, because we show that long-term deletion of *Insr* results in profound re-wiring of the β-cell transcriptome, the physiological changes can be the result of either direct or indirect action of *Insr* signaling. It should also be emphasized that, while we have deleted Insr, insulin can signal through Igf1r and Igf2r, especially at the higher concentrations predicted in the pancreas. A further caveat is that the molecular mechanisms involved in insulin secretion may be different in mouse and human β-cells ^82^, although we note that direction of effect we surmise agrees with the majority of human and canine studies, indicating general agreement across species^7, 8, 49, 50^.

In conclusion, our work demonstrates a modulatory role for autocrine insulin negative feedback and the lack thereof (i.e. β-cell resistance) in insulin secretion, glucose homeostasis and body mass, which depend on the physiological context studied. We hope our studies help resolve longstanding and controversial questions about the local effect of insulin on β-cells, and lead to experimental and theoretical studies that incorporate Insr-mediated signaling in other islet cell types.

## Materials and Methods

### Bioinformatics

Human tissue-level proteome and transcriptome were downloaded from the ProteomicsDB resource (https://www.proteomicsdb.org)83 in 2019 and sorted by relative protein abundance in Microsoft Excel. Publicly available human islet scRNAseq data were acquired from the panc8.SeuratData package and the SCTransform pipeline was followed to integrate the studies^84^. Expression data were normalized using the Seurat::NormalizeData function with default parameters and visualized using the Seurat::RidgePlot and Seurat::UMAPPlot functions, all from the Seurat package in R^85^.

### Mouse model and husbandry

All animal protocols were approved by the University of British Columbia Animal Care Committee (Protocol A16-0022), the Institutional Animal Care and Use Committee of the University of Massachusetts Medical School (A-1991-17) and the University of Michigan Institutional Animal Care and use Committee, in accordance with national and international guidelines. Mice were housed at room temperature on a 12/12 light dark cycle at the UBC Modified Barrier Facility, unless otherwise indicated.

Whenever possible, we used both female and male mice for experiments, and at multiple ages. However, we could not always conduct the studies on both sexes side-by-side under the same conditions, and in some cases the group sizes are un-even between sexes and age-groups. Thus, although we can make confident statistically-backed conclusions about the role of *Insr* in both female and male mice, at the specific ages studied, our study was not designed to explicitly compare sex or age as biological variables.

*Ins1*^cre^ mice on a C57Bl/6 background (mix of N and J NNT alleles) were gifted to us by Jorge Ferrer^30^(now commercially available, Jax #026801). The *Insr*^f/f^ allele on a pure C57Bl/6J background (#006955) and the nuclear TdTomato-to-nuclear EGFP (nTnG) lineage tracing allele^32^ on a mostly SV129 background with at least 2 backcrosses to C57Bl/6J before arriving at our colony were obtained from Jax (#023035) (Bar Harbor, ME). We generated two parental strains to avoid Cre effects during pregnancy; *Ins1*^cre/wt^;*Insr*^f/wt^ male mice and *Ins1*^cre/wt^;nTnG^/+^ female mice. These two parental strains were crossed in order to generate full littermate insulin receptor knockout *Insr*^f/f^;*Ins1*^cre/wt^;nTnG^/-^ (β*Insr*^KO^) mice, partial insulin receptor knockout *Insr*^f/wt^;*Ins1*^cre/wt^;nTnG^/-^ mice mice (β*Insr*^HET^), and their control groups *Insr*^wt/wt^;*Ins1*^cre/wt^;nTnG^/-^ mice (3 alleles of insulin) and *Insr*^f/f^;*Ins1*^wt/wt^;nTnG^/-^ mice (4 alleles of insulin). *Insr*^f^, *Ins1*^cre^, and nTnG genotyping were done in accordance with Jax’s recommendations using a ProFlex PCR system (Thermo Fisher Scientific, Canada). NNT genotyping was done as described previously^86^. Master mix for genotyping included 0.5μM primers (Integrated DNA technologies), 2mM dNTPs (New England Biolabs, #N0447S), 0.5U DreamTaq DNA polymerase (Fisher Scientific, #FEREP0702). Agarose gels varied from 1-2.5% (FroggaBio, #A87-500G). The mouse genetic background should be considered mixed, but relatively fixed and largely C57Bl6/J.

In our studies, mice were fed 1 of 3 diets: either a chow diet (PicoLab Mouse Diet 20-5058); a low-fat diet (LFD; Research Diets D12450B) containing 20% of kcal protein, 10% of kcal fat, and 70% of kcal carbohydrate including 35% sucrose, or; a high-fat diet (HFD; Research Diets D12492) containing 20% of kcal protein, 60% of kcal fat, and 20% of kcal carbohydrate including 10% sucrose.

MIP^cre^ mice were generously obtained from Dr Dempsey^87^. The CAG-YFP reporter transgenic animals were from Jax (#011107). All animals were males that were intraperitoneally injected at 8 weeks of age for 5 consecutive days with tamoxifen (Sigma, T5648) freshly dissolved in corn oil (Sigma, C8267) with 3 injections at 200 mg/kg over a 1-week period.

### Comparison of control genotypes

Before conducting our main study, we did a pilot experiment to determine whether the *Ins1*^cre^ knock-in mice had any phenotype on their own under both low fat and high fat diets, and we tracked both ‘control’ genotypes for the majority of our studies. Although *Insr*^wt/wt^;*Ins1*^cre/wt^;nTnG and *Insr*^f/f^;Ins1^wt/wt^:nTnG control mice exhibited generally similar phenotypes, we observed key differences that reinforced the rationale for using controls containing Cre and lacking 1 allele of *Ins1*, matching the experimental genotypes. For example, male HFD-fed *Insr*^f/f^;*Ins1*^wt/wt^;nTnG mice showed significantly higher levels of plasma proinsulin in comparison to *Insr*^wt/wt^;*Ins1*^cre/wt^;nTnG mice at 16 and 28 weeks of age (Fig. S7). At several ages, both LFD and HFD fed female *Insr*^f/f^;*Ins1*^wt/wt^;nTnG^/-^ mice exhibited trends toward slightly improved glucose tolerance (Fig. S8), most likely due to one extra allele of insulin, in comparison to *Insr*^wt/wt^;*Ins1*^cre/wt^;nTnG mice. Insulin sensitivity was generally similar, although not identical (Fig. S9). Longitudinal tracking of body weight revealed a consistent tendency for mice with a full complement of insulin gene alleles to be heavier than mice in which 1 allele of *Ins1* had been replaced with Cre. With the statistical power we had available, female HFD-fed *Insr*^f/f^;*Ins1*^wt/wt^;nTnG mice had significantly increased body mass at 11 and 16 weeks of age in comparison to *Insr*^wt/wt^;*Ins1*^cre/wt^;nTnG mice (Fig. S10). Once we had established the effects of the *Ins1*^cre^ allele on its own, we used a breeding strategy ensuring that all pups were born with 3 insulin alleles to control for any effects of reduced insulin gene dosage (See Fig. 1C). This strategy gave us cohorts of: *Insr*^wt/wt^;*Ins1*^cre/wt^;nTnG^/-^ (β*Insr*^KO^) control mice, *Insr*^f/wt^;*Ins1*^cre/wt^;nTnG^/-^ (β*Insr*^KO^) β-cell specific *Insr* heterozygous knockout mice, and *Insr*^f/f^;*Ins1*^cre/wt^;nTnG^/-^ (β*Insr*^KO^) β-cell specific *Insr* complete knockout mice. In some studies, the nTnG allele was not present (see Figure legends).

### Islet isolation and dispersion

Mouse islet isolations were conducted by ductal inflation and incubation with collagenase, followed by filtration and hand-picking as in our previous studies and following a protocol adapted from Salvalaggio et al. ^13, 15, 16, 88^. 24h post islets isolations, islets were washed (x4) (Ca/Mg-Free Minimal Essential Medium for suspension cultures, Cellgro #15-015-CV), followed by gentle trypsinization (0.01%), and resuspended in RPMI 1640 (Thermo Fisher Scientific #11875-093), 10%FBS, 1%PS). Cells were seeded either on glass cover slips or in 96-well plates according to the experimental procedure (see below).

### Immunoblotting

50 islets per sample were washed in PBS twice and then lysed and sonicated in RIPA buffer (10mM HEPES, 50mM β-glycerol phosphate, 1% Triton X-100) supplemented with complete mini protease inhibitor cocktail (Roche, Laval, QC) and phosphatase inhibitors (2mM EGTA, 70mM NaCl, 347 1mM Na3VO4, and 1mM NaF). Protein concentration was measured using Micro BCA Protein Assay Kit (ThermoFischer Scientific). 10 μg of total protein for each sample was separated by SDS-PAGE and transferred to Immun-Blot PVDF membrane (Bio-Rad Laboratories). Subsequently, membranes were blocked in I-Block (ThermoFischer Scientific) and probed with primary antibodies (Table 1 in Supplementary information) targeting insr-β subunit (1:1000, CST #3020S), ERK1/2 (1:1000, CST #4695), p-ERK1/2 (Thr202/Tyr204) (1:1000, CST #4370), AKT (1:1000, CST #9272), p-AKT (Thr308) (1:1000, CST #9275), ACTB (Novus Biologicals, NB600-501). Protein detection was performed by the use of the HRP-conjugated secondary antibodies: anti-rabbit (CST #7074) or anti-mouse (CST #7076) and Immobilon Forte Western HRP substrate (Millipore Sigma). Protein band intensity on exposed film was measured with Adobe Photoshop software.

### Targeted gene expression analysis

Tissue samples were kept frozen during grinding using mortals and pestles. cDNA was synthesized using qScript^TM^ cDNA synthesis kits (QuantaBio, #95047-500) following RNA was isolated from 50- 100mg of sample using RNeasy mini kits (Qiagen, #74106) according to manufacturer’s recommendations. qPCR was performed in 15ul reaction volumes using a CFX384 Touch Real-Time PCR Detection System (BioRad). Primer sequences: *Insr* forward 5’- 376 TTTGTCATGGATGGAGGCTA-3’ and *Insr* reverse 5’-CCTCATCTTGGGGTTGAACT-3’. *Hprt* forward 5’-TCAGTCAACGGGGGACATAAA-3’ and *hprt* reverse 5’-GGGGCTGT 379 ACTGCTTAACCAG-3’. *Insr* expression data were analyzed using the 2^-ΔΔ*CT*^ method using *Hprt* as a housekeeping gene. ΔCq=Cq(*Insr*)-Cq(*Hprt*) followed by normalization of the ΔCq(exp) to the mean of the *Hprt* expression in liver.

### RNA sequencing

To generate transcriptomic data from β-cells lacking *Insr* and littermate controls, groups of 50 islets were dispersed using mild trypsin digest according to our standard protocol^13, 44^, and FACS purified based on the GFP-positivity of the *Ins1*^Cre^-induced nTnG allele. RNA isolation and library preparation were conducted in accordance with the *SMART seq 2* protocol^89^. Sequencing of 100 GFP-positive β- cells was performed at the UBC Sequencing and Bioinformatics Consortium using Illumina NextSeq 500 with paired-end 75 bp reads. The number of read pairs per sample ranged from 4 million to 42 million, with a median of 18 million. Samples high glucagon reads (>10,000 CPM) were omitted, as presumed β-cell purification failures. Raw counts of gene reads were quantified by Kallisto^90^ and filtered for low count by only keeping genes with at least 5 counts in more than 10 samples. Using DESeq2 package^91^, gene counts were then normalized by variance stabilizing transformation and analyzed for differential expression using Benjamini–Hochberg adjusted p value < 0.05 as the cut-off. Similar results were found when multiple analysis pipelines were applied and their results combined^92^.

### Islet metabolism and oxygen consumption analysis

The Seahorse XF Cell Mito Stress Test kit (cat#103015-100) was used to measure oxygen consumption rate (OCR) in dispersed mouse islets using an Agilent Seahorse XF96 Analyzer (Seahorse Bioscience, North Billerica, MA). Dispersed islet cells were seeded at a density of 40,000 cells/well in XF culture microplates. After seeding (48h), the Seahorse XF Sensor Cartridge was hydrated in 180μL of Seahorse XF Calibrant Solution (cat#100840-000) added to each well of the XF Utility Plate (cat#102416-100). The hydrated cartridge was kept in a non-CO2 incubator at 37C° for 24h to remove CO2 from media. To pre-equilibrate the cells, 180 μL of Seahorse XF base medium (minimal DMEM) containing 10 mM glucose, 4 mM L-glutamine, and 2 mM sodium pyruvate was added to each well of the culture plate 1 h prior to the run and was also present during extracellular flux measurements. Mitochondrial respiration was analyzed by sequential injections of modulators including oligomycin (2 µM) used to block ATP synthase, carbonyl-cyanide-4-(trifluoromethoxy) phenylhydrazone (FCCP 3 µM) to activate uncoupling of inner mitochondrial membrane allowing maximum electron flux through the electron transport chain, and finally a mix of rotenone (1 µM) and antimycin A (1 µM) to inhibit complexes I and III, respectively. Mitochondrial proton leak, ATP-linked oxygen consumption and non-mitochondrial respiration were calculated based on the resulting respiratory profile, as described^93^.

### Patch-clamp electrophysiology

Islets from 16-week-old chow-fed male and female mice were isolated at UBC. 100-300 islets from each mouse shipped in a blinded manner overnight to the University of Alberta in RPMI (Invitrogen, 11875) with 10% FBS (Invitrogen #12483020), and 1% penicillin-streptomycin (Thermo Fisher, #15070063). Islets were dissociated into single cells using StemPro Accutase (Thermo Fisher Scientific, Cat# A11105-01) one day after receiving the islets. Dispersed cells were cultured in RPMI-1640 containing 11.1 mM glucose with 10% FBS and 100 U/ml penicillin/streptomycin for up to 2 days.

Membrane potential and current measurements were collected using a HEKA EPC10 amplifier and PatchMaster Software (HEKA Instruments Inc, Lambrecht/Pfalz, Germany) in either the current- or voltage-clamp mode in the perforated patch-clamp configuration. All the measures were done in a heated chamber (32–35°C). Membrane potential measurement was performed with patch pipettes pulled from thick-walled borosilicate glass tubes (Sutter Instrument), with resistances of 8–10 MΩ when filled with 76 mM K2SO4, 10 mM KCl, 10 mM NaCl, 1 mM MgCl2 and 5 mM Hepes (pH 7.25 with KOH), and back-filled with 0.24 mg/ml amphotericin B (Sigma, cat# a9528). The extracellular solution consisted of 140 mM NaCl, 3.6 mM KCl, 1.5 mM CaCl2, 0.5 mM MgSO4, 10 mM Hepes, 0.5 mM NaH2PO4, 5 mM NaHCO3, 5 mM glucose (pH 7.3 with NaOH). Membrane potential was measured with 5 mM G starting from the beginning, for 5 min, then changed to 1 mM G for 4-5 min, then changed to 10 mM G for 8-10 min, finally changed back to 5 mM G. KATP currents and reversal potential were recorded during and after membrane potential measurement on each cell. β-cells were distinguished by characteristic differences in the voltage-dependent inactivation of Na^+^ channel currents^94^.

Measurement of voltage-dependent exocytosis was performed in the whole-cell configuration. Before the start of whole-cell patch clamping, media were changed to bath solution containing (in mM): 118 NaCl, 20 Tetraethylammonium-Cl, 5.6 KCl, 1.2 MgCl2, 2.6 CaCl2, 5 HEPES, and 5 glucose (pH 7.4 with NaOH). For whole-cell patch-clamping, fire polished thin-walled borosilicate pipettes coated with Sylgard (3-5 MOhm), contained an intracellular solution with (in mM): 125 Cs-glutamate, 10 CsCl, 10 NaCl, 1 MgCl2, 0.05 EGTA, 5 HEPES, 0.1 cAMP, and 3 MgATP (pH 7.15 with CsOH). Quality control was assessed by the stability of seal (>10 GOhm) and access resistance (<15 MOhm) over the course of the experiment. Data were analysed using FitMaster (HEKA Instruments Inc) and Prism 6.0h (GraphPad Software Inc., San Diego, CA).

### Calcium imaging and analysis

Two days following cell seeding on glass coverslips, adherent islet cells were loaded with 5μM of the acetoxymethyl (AM) ester form of the calcium indicator Fura-2 (Thermo Fisher Scientific #F1221) for 30min. Islet cells were perifused at 1mL/min for 45min prior to experimental procedure to ensure washout of excess FURA2. During experiments, cells were mounted on a temperature-controlled stage and held at 37°C on a Zeiss Axiovert 200M inverted microscope equipped with a FLUAR 20× objective (Carl Zeiss, Thornwood, NY), while perifused with Krebs-Ringer (KRB) solution (144mM NaCl, 5.5mM KCL, 1mM MgCl2, 2mM CaCl2, 20mM HEPES) of various glucose concentrations as indicated in figures. Ca^2+^ traces were analyzed automatically, as follows. Taking a similar approach to that described previously^34^, 8 features were extracted from the Traces (Fig S2). Peaks during each phase were identified as local maxima reaching a value with a percent difference above the median baseline level greater than 20%. P-values for calcium analysis were generated using ANOVA with correction for multiple comparisons performed using Tukey’s method. Figures were generated using the ggplot2 package in R (Wickham2016). Code used to analyze data and generate figures are available upon request.

### Analysis of total protein synthesis rate

For the purpose of pulse labeling of newly translated proteins, 50 isolated islets were incubated in complete RPMI media without cysteine and methionine (MP Biomedicals, #SKU 091646454) for 1hr. Subsequently media was supplemented with 250 μCi of [^35^S]-cysteine/methionine mixture (PerkinElmer, NEG772002MC) and islets were incubated under normal conditions for 30 min. Islets were then lysed and proteins separated by SDS-gel electrophoresis as described above. Gels were fixed for 30 min in 50% (v/v) ethanol in water with 10% (v/v) acetic acid, dried in gel dryer (Bio-Rad model 583) and then exposed to storage phosphor screen (GE Healthcare) overnight. Screens were imaged and digitised using Typhoon FLA 9000 biomolecular imager (GE Healthcare). Protein bands intensity was quantified with Adobe Photoshop software.

### Dynamic insulin secretion perifusion analysis

For perifusion experiments, islets from 16week old chow-fed male and female mice were isolated using collagenase, filtration and hand-picking as previously described^95^. Our standard approach compared the insulin response to both 20mM and 10mM glucose stimulation as well as direct depolarization with 30 mM KCl. More specifically, 150 hand-picked islets per column were perifused (0.4 mL/min) with 3 mM glucose KRB solution containing (in mM) 129 NaCl, 4.8KCL, 1.2 MgSO4•7H2O, 1.2 KH2PO4, 2.5 CaCl2, NaHCO3, 10 HEPES, as well as 0.5% BSA (Sigma # A7030) for 60 min to equilibrate the islets to the KRB and flow rate, and then treated as indicated. Samples were analyzed using a rat insulin radioimmunoassay that has 100% cross-reactivity for mouse insulin (Millipore-Sigma #ri-13k). Insulin content was measured after acid-ethanol extraction using an insulin ELISA (Stellux Rodent Insulin ELISA, Alpco #80-INSMR-CH10).

### Hyperglycemic clamps to assess glucose-stimulated insulin secretion in vivo

*In vivo* hyperglycemic clamp experiments were performed at the National Mouse Metabolic Phenotyping Center (MMPC) at UMass Medical School. Body composition analysis was conducted by noninvasively measuring whole body fat mass and lean mass using ^1^H-MRS (Echo Medical Systems, Houston, TX). A survival surgery was performed at 5–6 days before hyperglycemic clamp experiments to establish an indwelling catheter in the jugular vein. On the day of experiment, mice were fasted overnight (∼15h), and a 2-h hyperglycemic clamp was conducted in awake mice by intravenously infusing 20% dextrose to rapidly raise and maintain plasma glucose levels at ∼19 mM^96^. Blood samples were taken at 10∼20 min intervals to measure plasma insulin levels during hyperglycemic clamps.

### Intravenous 4-day glucose infusion

Mice were bred and genotyped at University of British Columbia and shipped at 5 weeks of age to the Division of Diabetes, Department of Medicine, University of Massachusetts Medical School, USA. In a blinded manner, glucose infusions were performed as described^97^. Jugular vein catheters were placed in 9-12-week-old male and female mice with blinded genotypes. From postoperative recovery through euthanasia mice were unrestrained and were maintained on a 12-h light/dark cycle, with access to 2.2 g diet (to ensure isocaloric intake across all mice) and water. After 2 days of recovery, mice received continuous 4-day intravenous infusions of 50% dextrose (Baxter) containing 500ug/mL BrdU. Tail blood was sampled for plasma insulin, glucagon and blood glucose at Day 0, 1, 2 and 4. Blood glucose was measured using ReliOn glucometer (Walmart), glucagon was measured using mouse Glucagon ELISA (Mercodia 10-1281-01), and plasma insulin was measured using mouse Insulin ELISA kit (Mercodia 10-1247-01). Mice were euthanized at the end of the experiment and pancreas and duodenum were harvested for histology. Tissues were fixed for 5 h in 10% formalin and then stored in 1X PBS until processing, paraffin embedding and sectioning. Images were acquired using a NIKON fully motorized for Phase and Fluorescence Ti-E microscope. Images were taken of at least 10 randomly selected islets, all four channels at the same time. To generate RGB images, channels were inserted to show Insulin-BrdU-DAPI, Glucagon-BrdU-DAPI or Insulin-Glucagon-DAPI. To generate yellow-magenta-white images to accommodate colorblind viewers, new files were generated in Adobe Photoshop in which original channel data were displayed in multiple channels using the merge function (e.g. to change green to yellow, green channel data were added to both green and red channels; for more detailed information please contact LCA). Cells were counted using Cell profiler automated counting software from Broad Institute (Cambridge, MA); all counts were manually checked.

### β-Cell mass and immunohistochemistry

Pancreata were perfused, then fixed for 24h with 4% paraformaldehyde, and then washed twice with 70% ethanol prior to paraffin embedding and sectioning (5 μm) to obtain 5 different regions of the pancreas (100 μm apart) by WAXit Inc. (Vancouver, Canada). Paraffin was removed by 5 min xylene incubation steps. Sections were rehydrated in decreasing concentrations of ethanol and rinsed with water and PBS. Epitope retrieval was done either by immersing samples 10 mM citrate buffer, pH 6.0 for 15min at 95°C, or by transferring sections to prewarmed 1N HCl for 25 min at 37°C. Samples were washed with PBS twice and were either blocked for 10 min at room temperature (Dako protein block #X0909), or with goat block (GB) with Triton X-100 (10% BSA + 5% Goat Serum with 0.5% Triton X- 100) for 1-4 h at room temperature. Samples were incubated overnight at 4°C in primary antibodies targeting anti-insulin (1:200 Abcam #Ab7872), anti-glucagon (1:100 Cell Signaling Technologies, #2760S), anti-BrdU (1:250, Abcam ab6326), anti-GLUT2 (1:1000, Milipore, #07-1402). Following 3 PBS washes (5 min each), samples were incubated for 30min or 1h at room temperature in secondary antibodies in a light-deprived humid chamber. Secondary antibodies applied were anti-rabbit Alexa Fluor-488 (1:200, Invitrogen, # A-11008), anti-rabbit Alexa-488 (1:200, Invitrogen, #A11034), anti-rat Alexa-594 (1:200, Invitrogen, #A11007), anti-guinea pig Alexa-647 (1:200, Invitrogen, #A21450), anti-guinea pig Alexa-594 (1:200, Invitrogen #A-11076). Samples were mounted with either VECTASHIELD Hard Set Mounting Medium (Vector labs, # H-1500) or Fluoroshield both containing DAPI (Sigma-Aldrich, #F6182-20ML) following an additional three washes in PBS (10 min each). For β-cell area quantification, whole pancreas sections were imaged using an ImageXpress^MICRO^ using a 10x (NA 0.3) objective and analyzed using the MetaXpress software (Molecular Devices, San Jose, CA, USA). Beta cell area was calculated as insulin positive area normalized to the entire pancreas of each section. The mean of five sections from 5 regions of the pancreas were quantified. For other immunofluorescence analysis of fixed tissue, we used a Zeiss 200M microscope using 20x air objective (NA 0.75), NIKON fully motorized for Phase and Fluorescence Ti-E microscope. For live cell imaging for recombination validation, islets from *Ins1*^cre/wt^:nTnG mice were incubated with CellMask™ Deep Red Plasma membrane stain (Thermo Fisher #C10046) using a Leica confocal microscope.

For β-cell mass analysis of *Insr* knockouts using MIP-Cre, β-cell mass determination was performed by intensity thresholding using the Fiji 2.1.0-/1.53c image analysis package, after fluorescence immunostaining for insulin on 5 independent sections per animal, selected at random throughout the pancreas as previously described. The proportion of insulin positive staining to total area was them multiplied by the pancreatic weight to derive the β-cell mass. The data was subsequently analyzed using a non-parametric statistical test (Mann-Whiney) in Prism 8.4.3 (GraphPad, San Diego, CA).

### Blood collection and in vivo analysis of glucose homeostasis and insulin secretion

Tail blood was collected for blood glucose measurements using a glucometer (OneTouch Ultra 2 meter, Lifescan, Canada) for single time points as well as during glucose and insulin tolerance tests. Mice were fasted for 4h or 16h during single timepoints and for 6h during glucose and insulin tolerance tests, as well as glucose stimulated insulin secretion tests. The *i.p.* glucose dose was 2g/kg unless otherwise specified. The *i.p.* Humalog (Eli Lilly and Co) insulin dose was 0.75U unless otherwise indicated. 1-2 days prior to femoral blood collection the experimental mice were tube handled and shaved. Femoral blood was collected for single timepoints, as well as for measurements of *in vivo* glucose-stimulated insulin secretion after *i.p.* injection of 2g/kg glucose. Blood samples were kept on ice during collection, centrifuged at 2000rpm for 10min at 4°C and stored as plasma at -20°C. Plasma samples were analysed for insulin (Stellux Chemi Rodent Insulin ELISA, Alpco #80-INSMR-CH10), proinsulin (Alpco #80-PINMS-E01), C-peptide (Alpco #80-CPTMS-E01), glucagon (Mercodia, #10- 1281-01). Measurements were performed on a Spark plate reader (TECAN), and analysed using (GraphPad Software Inc., San Diego, CA).

### Mathematical modelling

We used modified versions of the Topp model (Topp et al., 2000) to simulate glucose, insulin, and, in certain instances, β-cell dynamics (see equations and detailed process in Fig. 6). To capture the proposed insulin-receptor mediated negative feedback on insulin secretion, we introduced an inhibition factor, 1/(1 + *Sβ I*), multiplying the insulin secretion term, parametrized by Sβ, the β-cell Insr-specific insulin sensitivity. The other structural modification to the Topp model that we made was the removal of the β-cell mass equation when modelling glucose tolerance tests, justified by the slow nature of changes to β-cell mass compared to the fast changes in glucose and insulin levels during the test.

In glucose clamp conditions, a simple equation, derived in the Supplemental Material, relates Sβ to steady state insulin levels in wildtype and mutant mice: *Sβ* = (*Im* * - *Iwt* *) / (*Im* *)^2^, where *Iwt* * is the steady * state insulin level for wild type control mice, and *Im* is the equivalent for the β*Insr*KO mice. We assumed that, at each time point after the first one in the glucose-clamp insulin measurements, the blood insulin levels were hovering around steady state and all other parameters and variables were constant (Fig. S5). We used those insulin steady-state data points together with the simple equation to estimate Sβ for the male and female wild-type mice. The equation for Sβ is parameter free, independent of the β-cell mass steady state, and depends only on the data so no parameter nor β-cell mass estimates were required.

For *in silico* glucose tolerance tests, we varied the values of Sβ and SP to see their influence on glucose homeostasis. To set the parameter values in our model for this exploration, we started with the Topp model, originally parametrized for humans and gradually varied them with two simultaneous objectives: (1) ensure that a simulated glucose tolerance test roughly matched a typical wildtype female low-fat diet glucose tolerance test time series and (2) the steady state β-cell mass of our full GIβ model roughly matched a typical observed β-cell mass in these mice. The resulting parameter values are given in (Fig. 6). For *in silico* glucose tolerance tests, we then inserted the negative-feedback factor in the insulin secretion term (referred to here as *our full GIβ model or our fast GI model for the constant-β version*) and used the parameters values estimated as described above. We held the β-cell mass constant at the steady state β value of our full GIβ model, calculated using the clamp-data estimated Sβ value for wildtype female LFD mice and the SP value from the rough parameter estimation described above. For any individual tolerance test, we set Sβ and SP, calculated the steady state of our fast GI model and used the steady state insulin value as the initial condition for insulin and the steady state glucose value plus 20 mM as the initial condition for glucose. Fig. 6A shows sample *in silico* glucose tolerance tests for various values of Sβ, with SP fixed at 70. The area under the glucose curve (AUGC) for the glucose tolerance test was then calculated for a range of Sβ and SP values, generating a numerical map from Sβ and SP to AUGC. From the glucose tolerance tests in wildtype and mutant male mice and wildtype and mutant female mice, we calculated the experimental AUGC values at 4, 9, 21 and 39 weeks. Using the computed AUGC map and the 11-week Sβ values calculated from the glucose-clamp data, we estimated the SP value that would give the experimental AUGC.

### Statistics

Data are presented as mean +/- SEM, with individual data points from biological replicates, unless otherwise indicated. T-tests were performed when only 2 groups were compared at a single timepoint. One-way ANOVA was applied when three groups were compared at a single timepoint. Mixed effects model statistics were used for statistical analysis of GTT, ITT, GSIS experiments. Specifically, a fitted mixed model (Prism 8, Graph Pad), which allows for missing measurements and un-even group sizes, was applied when 2 or more groups were compared at multiple time points (e.g. 4h fasted blood glucose, insulin, proinsulin, c-peptide and body weight). We corrected for multiple comparisons within experiments, for example by using Bonferroni adjusted p values in the RNA sequencing analysis.

For glucose tolerance test data, we also used a Bayesian approach with multilevel regression modelling was used due to the high number of correlated experiments with repeated measurements performed on the animals^98^, in R version 4.1.0^99^. Missing values were multiply imputed using Amelia^100^ with an m parameter of 10 if not missing at random (e.g. samples above or below a limit of detection) using priors of the limit of detection. Generally, a linear mixed effects model was fitted with each animal treated as a random variable contributing to each measurement, and time treated as ordinal (minutes) or nominal (weeks). All Bayesian models were created in Stan computational framework accessed with the brms package ^101, 102^. Posteriors were checked for goodness of fit and convergence. Default parameters for chain length, max_treedepth, and adapt_delta were used with 8000 iterations. Weakly informative priors were specified. If no result is reported, the posterior probability did not exceed 95%. Bayes factors are reported and strength of evidence assessed as per convention^103^. In related figures, lines are the fitted values of the posterior distribution with shading indicating the 95% credible interval. Related figures were generated using ggplot2^104^. Related tables were generated using gt (https://CRAN.R-project.org/package=gt).

## Data availability

Raw RNAseq data are deposited in GEO (GSE159527)

## Author contributions

S.S. led project design/management, conducted experiments, analyzed data, and wrote manuscript

E. P. conducted experiments, analyzed data, and edited manuscript (protein synthesis, Westerns)

J. K. conducted experiments, analyzed data, and edited manuscript (ex vivo insulin secretion)

H. H. C. analyzed data, and edited manuscript (RNAseq analysis)

D. A. D. conducted experiments and analyzed data (in vivo glucose homeostasis)

X-Q. D. conducted experiments, analyzed data, and edited manuscript (electrophysiology)

R. B. S. conducted experiments, analyzed data, and edited manuscript (long-term hyperglycemia, proliferation)

L. E. conducted experiments, analyzed data, and edited manuscript (MIPCre mice)

C. E. analyzed data, and edited manuscript (Bioinformatics and calcium data analysis)

K. F. analyzed data, and edited manuscript (mathematical modelling)

S. A. M. M. conducted experiments and analyzed data (beta-cell area)

N. N. conducted experiments (imaging)

P. O. conducted experiments, analyzed data, and edited manuscript (Seahorse)

D. H. conducted experiments (mRNA expression qPCR)

X. H. conducted experiments (in vivo physiology)

H. L. conducted experiments (in vivo physiology)

H. M. conducted experiments, analyzed data, and edited manuscript (FACS, RNAseq prep)

J. W. conducted experiments (beta cell INSR western blots)

J. D. B. conducted experiments (liver INSR western blots)

H. L. N. conducted experiments (hyperglycemic clamps)

S. S. conducted experiments (hyperglycemic clamps)

A.B. conducted experiments (electrophysiology)

R.K. conducted experiments (electrophysiology)

B. G. conducted experiments (long-term hyperglycemia, proliferation)

C. C-M. conducted experiments, and edited the manuscript (MIPCre mice)

S. F. analyzed data (RNAseq analysis)

S. S. conducted experiments (RNAseq)

D. S. L. conducted experiments, analyzed data, and edited manuscript (Seahorse, nTnG validation)

C. N. supervised experiments, and edited manuscript (RNAseq)

E.J.R supervised experiments, and edited manuscript

E.N.C. supervised experiments, and edited manuscript (mathematical modeling)

J. K. supervised studies, and edited manuscript (Hyperglycemic clamps)

E. B-M. supervised studies, and edited manuscript (MIPCre mice)

L. A. supervised studies, and edited manuscript (long-term hyperglycemia, proliferation)

P. E. M. supervised studies, and edited manuscript (electrophysiology)

J. D. J. conceived the project, oversaw its execution, edited manuscript, and guarantees the work

## Acknowledgements

We thank Johnson lab members for valuable feedback and those who provided insight when data was presented at conferences. We thank Sarah Hulme, Charles Nieh, and our animal care staff for supporting our animal husbandry, Bernard Thorens and Jorge Ferrer for sharing the *Ins1*^cre^ knock-in mice, Eric Jan for allowing us to use his facilities to conduct S^35^ labeling. We thank Quin Wills for helpful discussion of the smartseq2 protocol, and Austin Taylor for discussion of genotyping.

## Funding

SS was supported by an Alfred Benzon Post-Doctoral fellowship. Work was primarily supported by a CIHR grant (133692) to JDJ. Work in Edmonton was supported by CIHR grant (148451) to PEM. Work in the Michigan Diabetes Research Center was supported by NIH (P30 DK020572). Work at U Mass Medical School was supported by NIH grants (5U2C-DK093000) to JKK and (R01DK114686) to LCA. CN is supported as a Tier 1 CRC.

## Conflict of Interest Statement

The authors report no conflicts of interest related to this work.

---

**Fig. S1.**
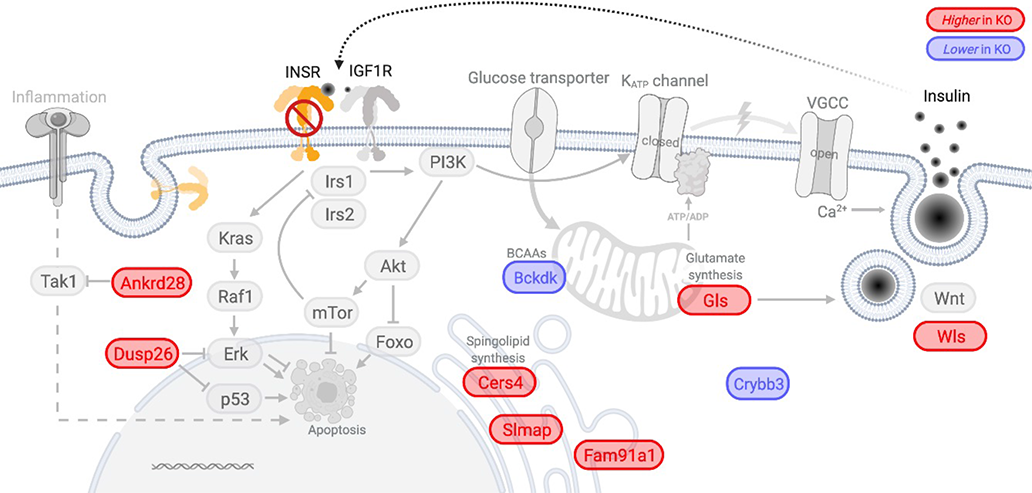
Possible roles of differentially expressed genes in beta-cells lacking Insr. The diagram depicts functions of genes differentially expressed between beta-cells of wildtype controls and beta-cell specific Insr KO mice (both sexes pooled).

**Fig. S2.**
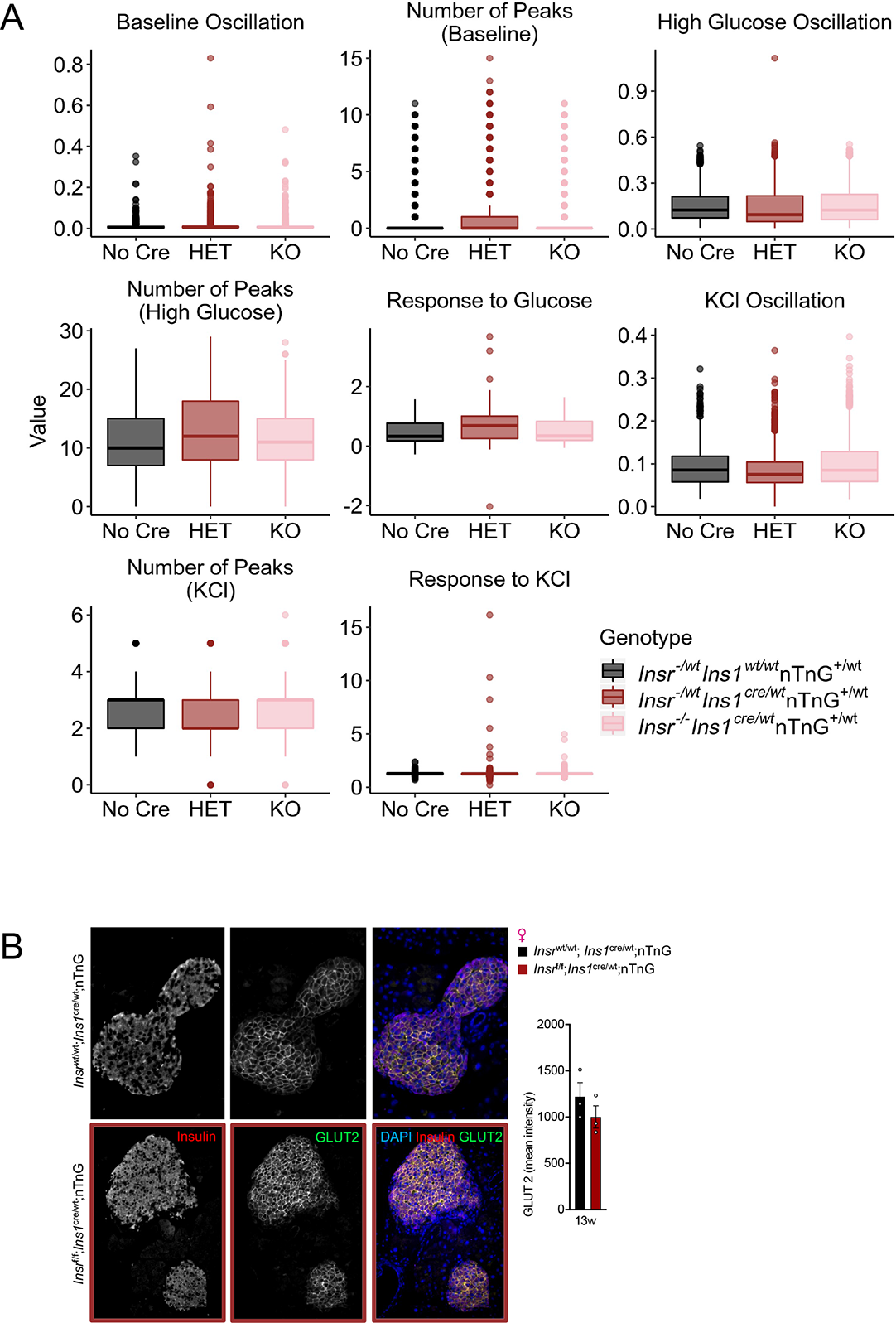
Additional quantification of dynamic Ca^2+^ responses. **(A)** Responses to glucose or KCl were defined as the median high glucose (15mM) or KCl (30mM) signal above baseline glucose (3mM), respectively, and normalized to the maximum response to KCl above baseline. High glucose-stimulated Ca^2+^ oscillations, Ca^2+^ oscillation in low glucose, and KCl-stimulated Ca^2+^ oscillation were defined as the median absolute deviation (MAD) during the high glucose, low glucose or KCl exposures, respectively, normalized to the maximum response to KCl above baseline. **(B)** Representative image and quantification of Slc2a2 (Glut2) in islet from sectioned pancreas from a LFD-fed 13 week old female control and β*Insr*^KO^ mice. Unpaired t-test.

**Fig. S3.**
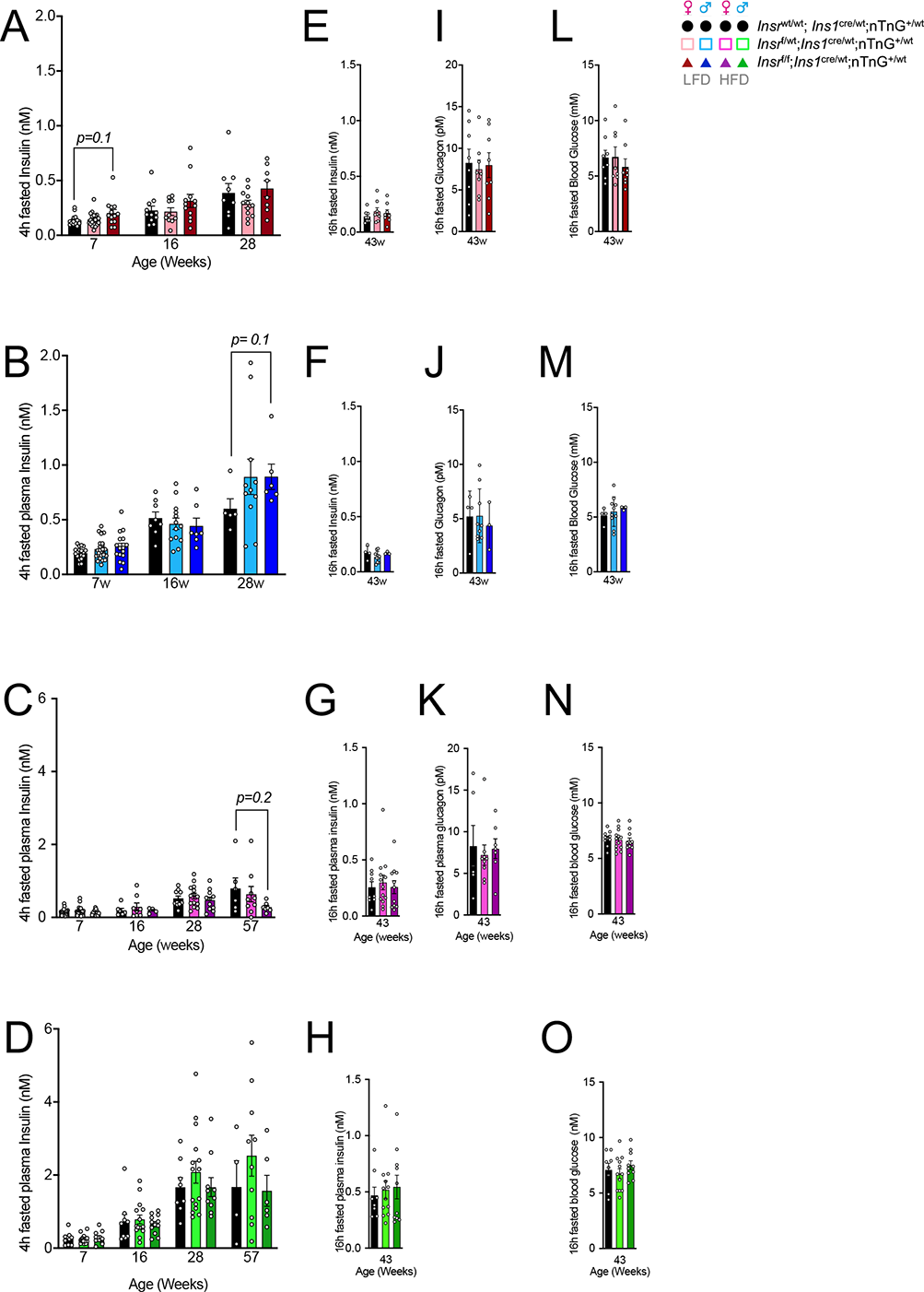
Fasting insulin, glucagon, and glucose levels. **(A-D)** Circulating plasma insulin levels after a 4-hour fast, **(E-H)** or a 16-hour fast; **(I-K)** plasma glucagon levels after a 16-hour fast **(L-O)** plasma glucose concentration after a 16-hour fast in control, β*Insr*^HET^, and β*Insr*^KO^ mice at multiple ages.

**Fig. S4.**
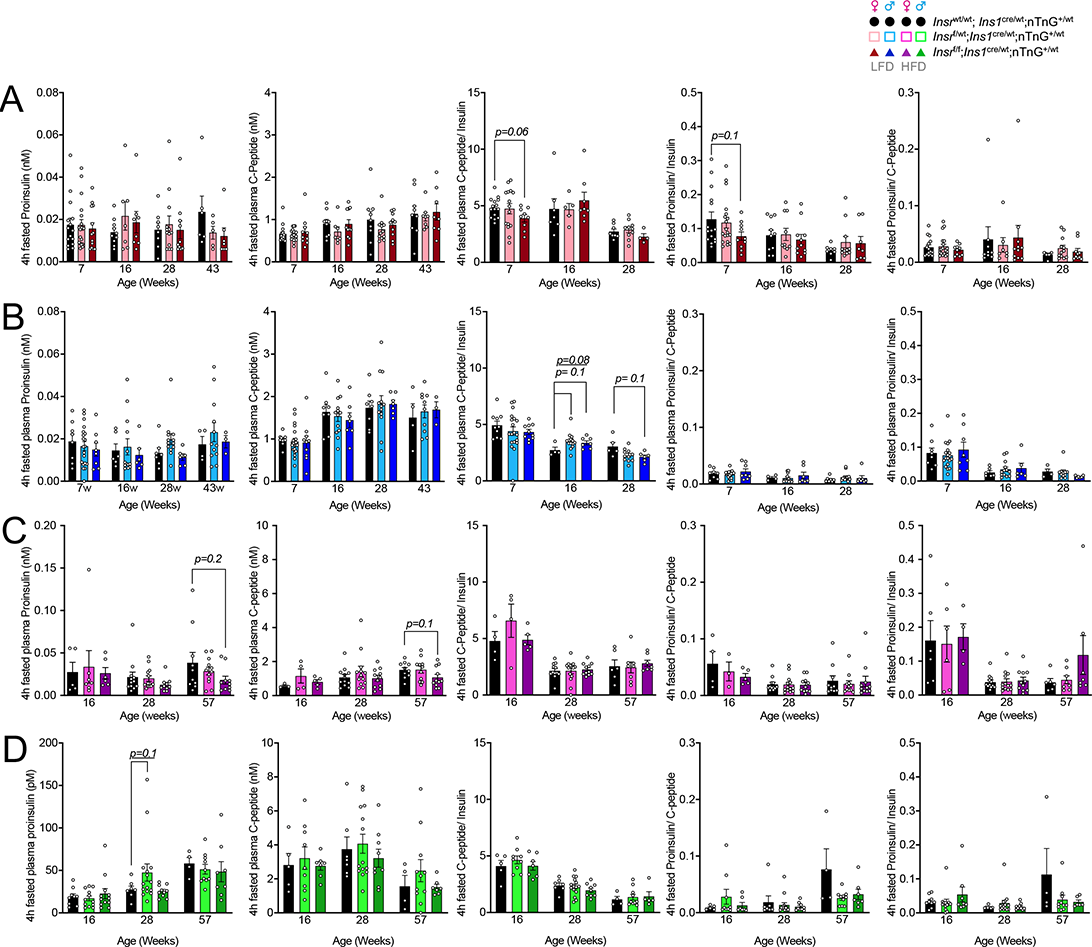
Fasting proinsulin and C-peptide. **(A-D)** Circulating plasma proinsulin and C-peptide levels after a 4-hour fast, and their associated ratios in control, β*Insr*^HET^, and β*Insr*^KO^ mice at multiple ages. Statistical analyses were done with a mixed model.

**Fig. S5.**
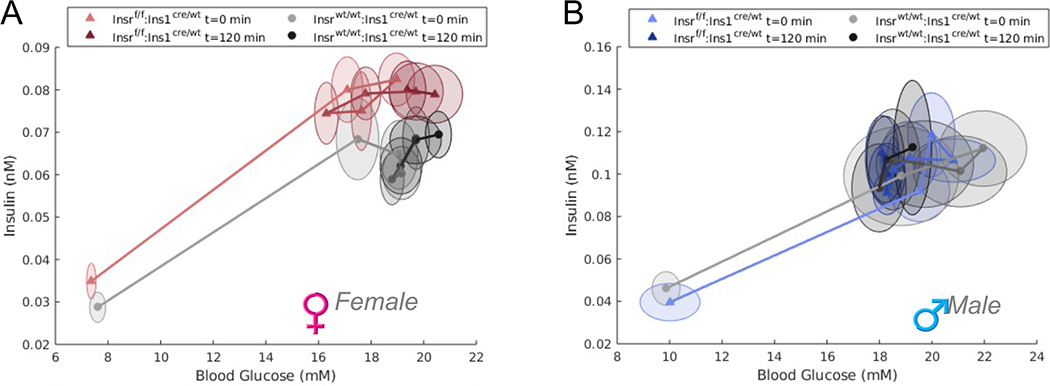
Mathematical modeling of hyperglycemic clamp data. **(A,B)** Relationship between insulin and glucose during the hyperglycemia clamps over time in female and male mice. Data from the clamp studies were used to define a beta-cell insulin sensitivity term that was included in a modified Topp model (see main text).

**Fig. S6.**
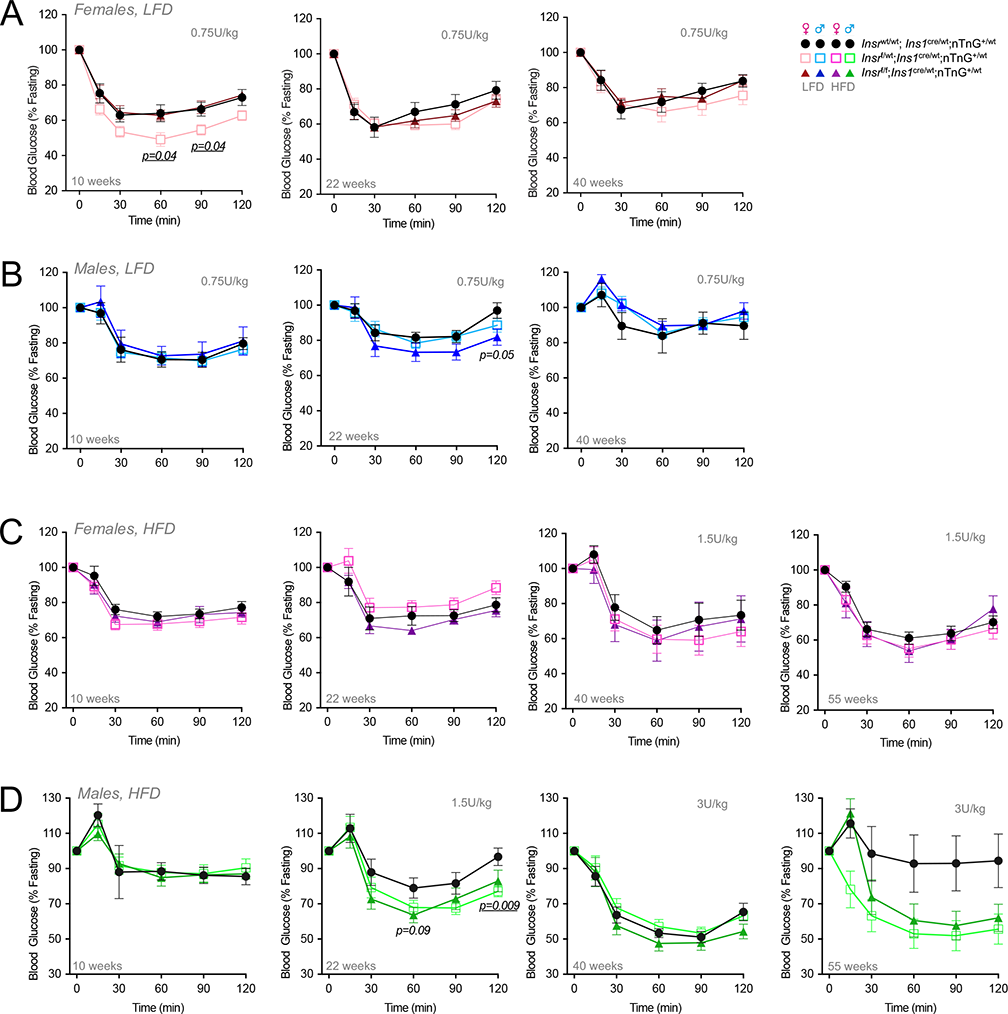
Insulin tolerance tests. **(A-D)** Insulin tolerance tests after a 6 hour fast in control, β*Insr*^HET^ and β*Insr*^KO^ mice fed LFD or HFD at multiple ages (n=5-26). Statistical analyses were done with repeated measures 2-way ANOVA. Doses are 0.75 U/kg unless otherwise shown.

**Fig. S7.**
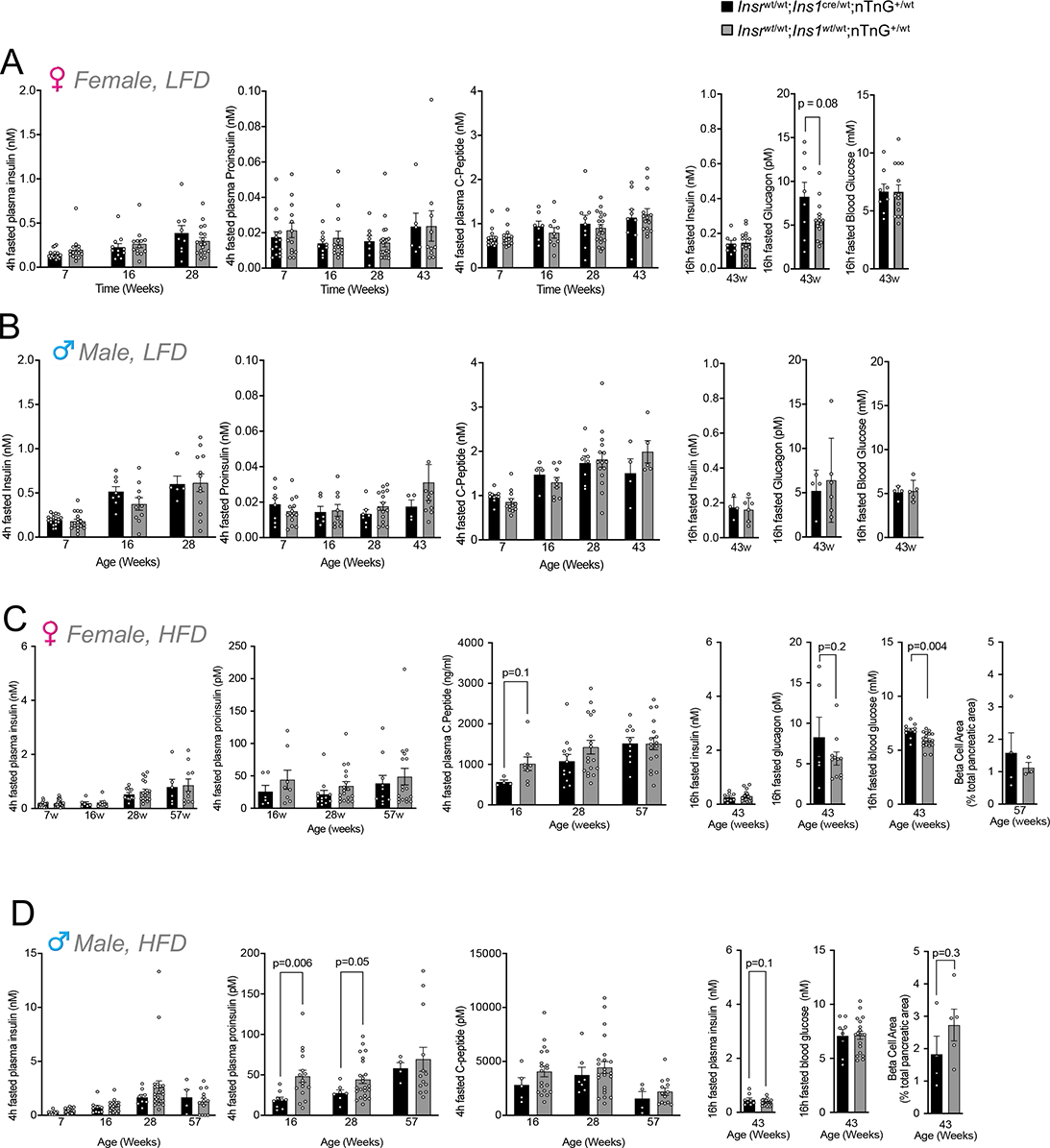
Fasting insulin, proinsulin, C-peptide and glucagon in control mice. **(A-D)** Circulating plasma insulin, proinsulin and C-peptide levels after a 4-hour fast, and their associated ratios as well as 16-hour fasted insulin, glucagon and blood glucose levels in *Insr*^wt/wt^;*Ins1*^Cre/wt^;nTnG^+/-^ (*black*) and *Insr*^wt/wt^;*Ins1^wt^*^/wt^;nTnG^+/-^ (*grey*) LFD-mice at multiple ages.

**Fig. S8.**
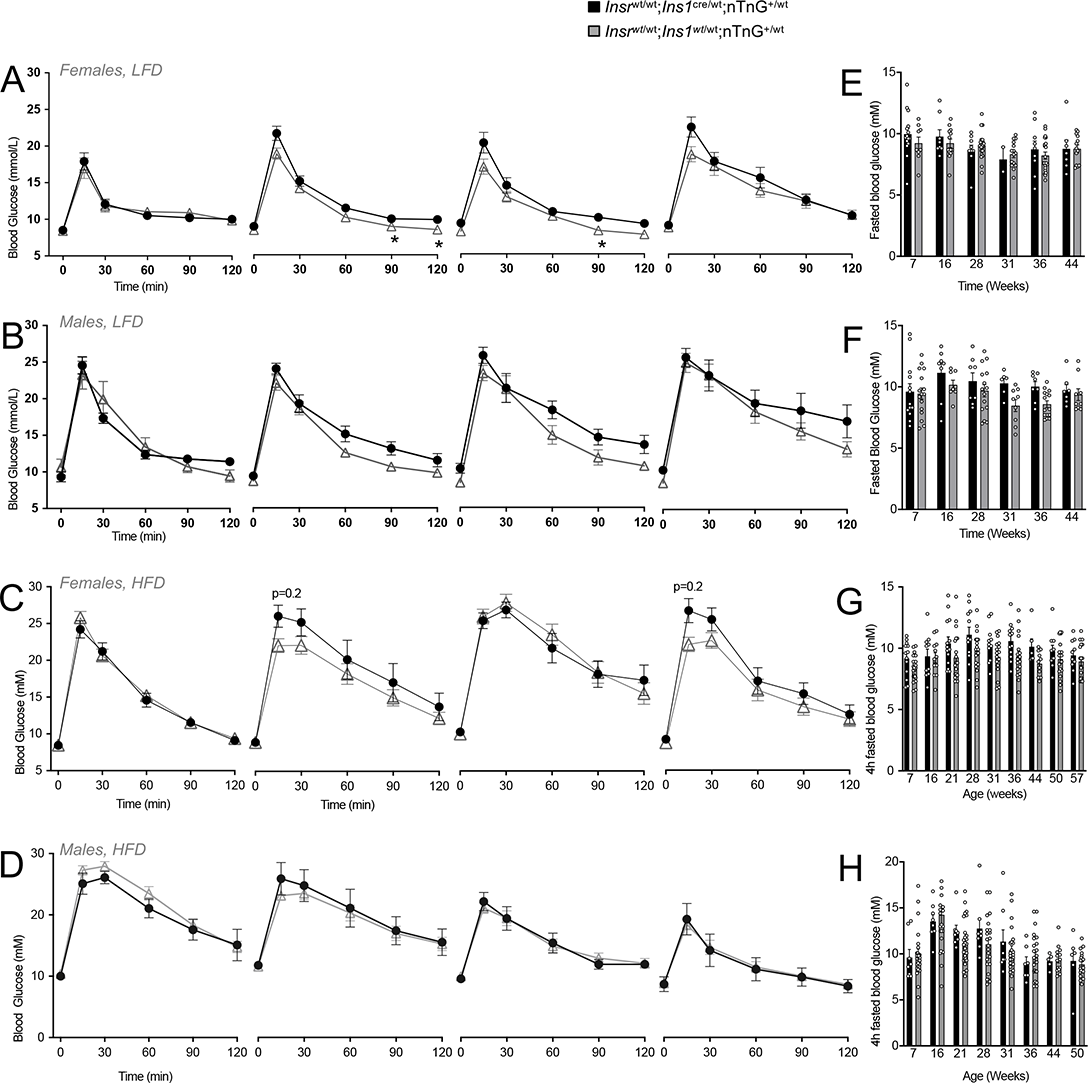
Glucose tolerance and fasting glucose comparison in controls. **(A-D)** Glucose tolerance tests after a 6 hour fast in female and male of *Insr*^wt/wt^;*Ins1*^Cre/wt^;nTnG^+/-^ (black) and *Insr*^wt/wt^;*Ins1^wt^*^/wt^;nTnG^+/-^ (grey) fed LFD or HFD at multiple ages (n=5-30). Statistical analyses were done with repeated measures 2-way ANOVA. **(E-H)** Blood glucose after a 4 hour fast.

**Fig. S9.**
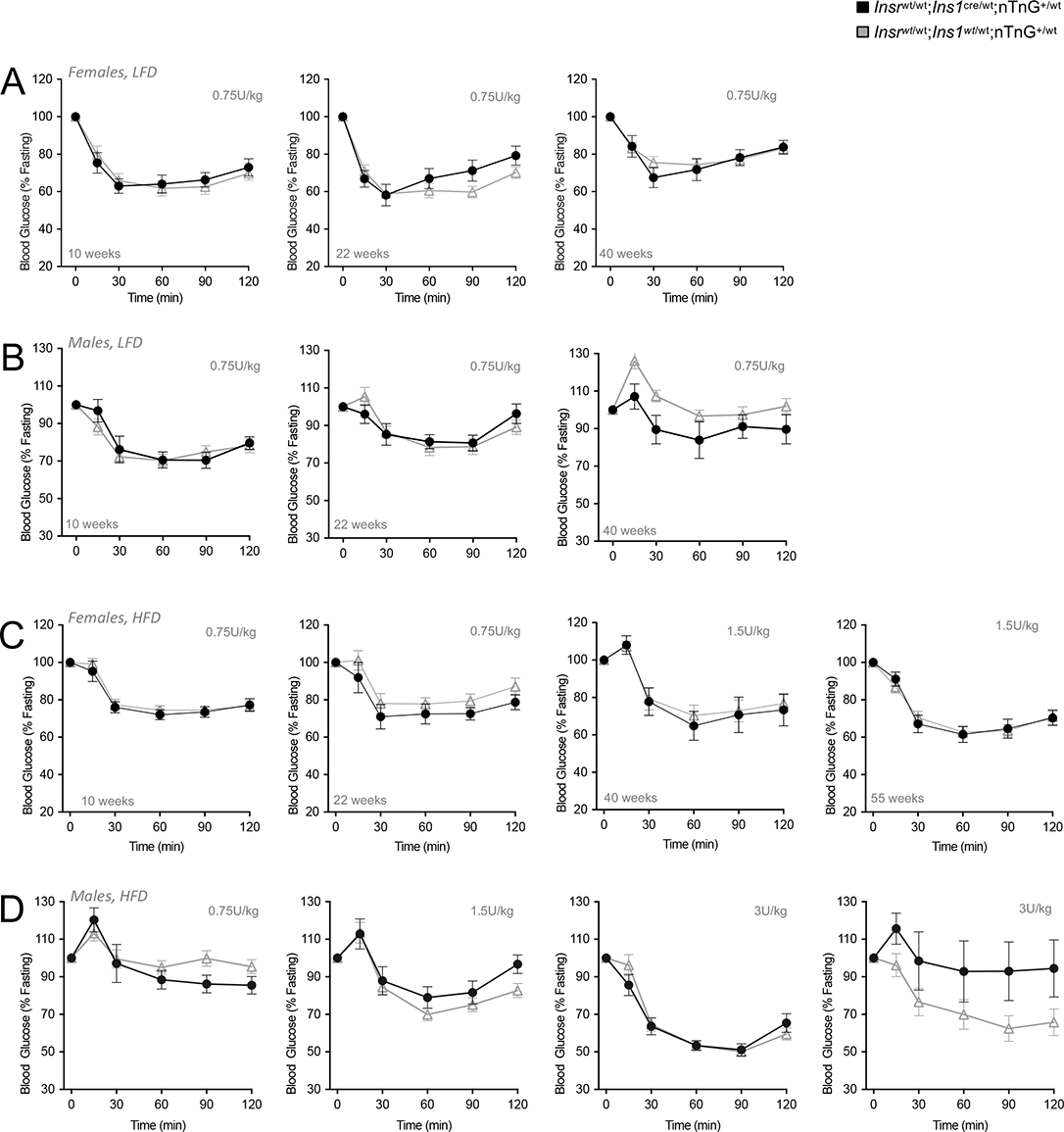
Insulin tolerance tests in control mice. **(A-D)** Insulin tolerance after a 6 hour fast in female and male of *Insr*^wt/wt^;*Ins1*^Cre/wt^;nTnG^+/-^ (black) and *Insr*^wt/wt^;*Ins1^wt^*^/wt^;nTnG^+/-^ (grey) fed LFD or HFD at multiple ages (n=5-30). Statistical analysis were done with repeated measures 2-way ANOVA. Doses are 0.75 U/kg unless otherwise shown.

**Fig. S10.**
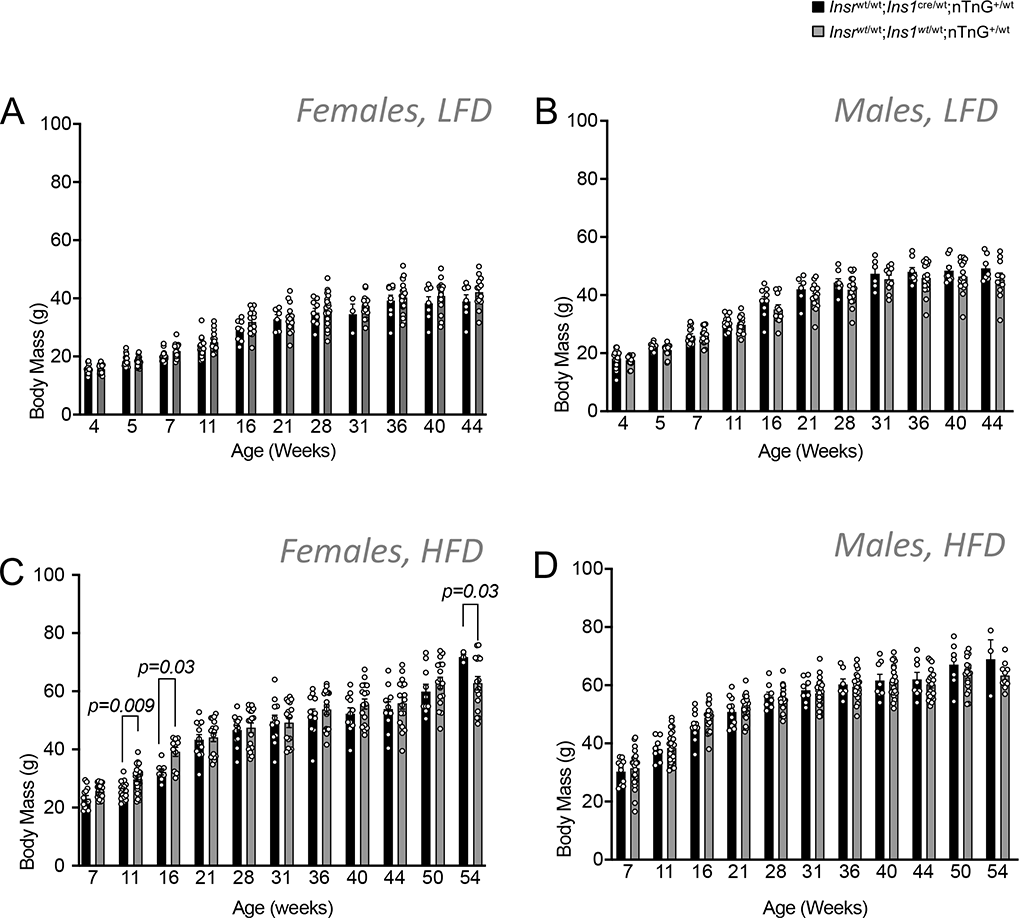
Body weight in control mice. **(A-D)** Body weight in female and male of *Insr*^wt/wt^;*Ins1*^Cre/wt^;nTnG^+/-^ (black) and *Insr*^wt/wt^;*Ins1^wt^*^/wt^;nTnG^+/-^ (grey) fed LFD or HFD at multiple ages. Statistical analysis with mixed effect model.

**Table S1.**
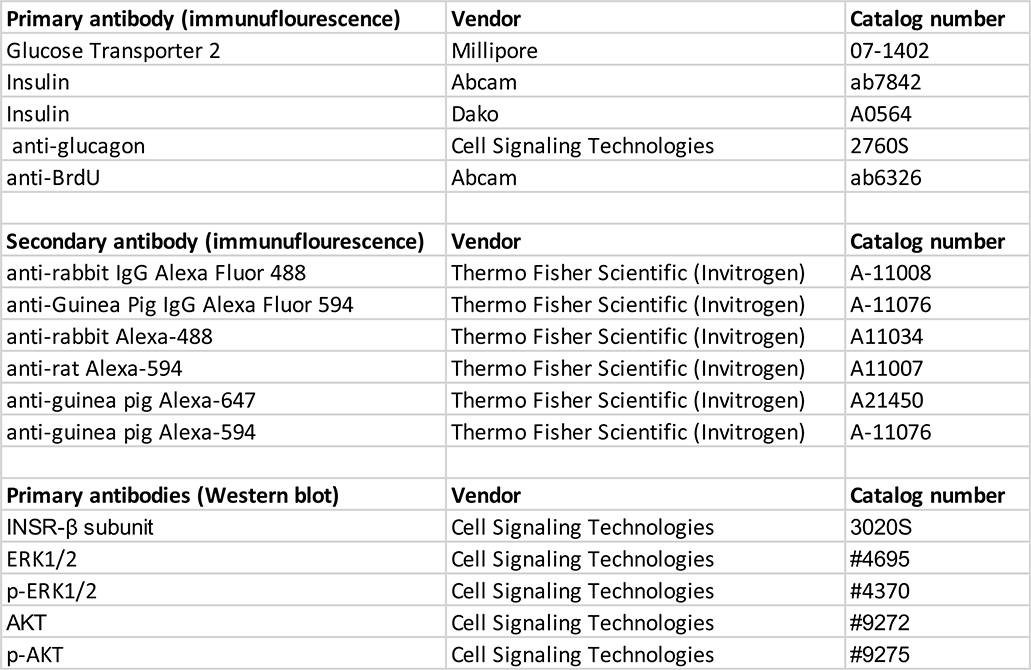
List of antibodies.

## Notes

### Competing Interest Statement

The authors have declared no competing interest.

### Summary of Updates

Minor revisions requested in peer review.

